# Demographic history and the efficacy of selection in the globally invasive mosquito *Aedes aegypti*

**DOI:** 10.1101/2024.03.07.584008

**Authors:** Tyler V. Kent, Daniel R. Schrider, Daniel R. Matute

**Author notes:** Address correspondence: Daniel R. Schrider or Daniel R. Matute.

## Abstract

*Aedes aegypti* is the main vector species of yellow fever, dengue, zika and chikungunya. The species is originally from Africa but has experienced a spectacular expansion in its geographic range to a large swath of the world, the demographic effects of which have remained largely understudied. In this report, we examine whole-genome sequences from 6 countries in Africa, North America, and South America to investigate the demographic history of the spread of *Ae. aegypti* into the Americas its impact on genomic diversity. In the Americas, we observe patterns of strong population structure consistent with relatively low (but probably non-zero) levels of gene flow but occasional long-range dispersal and/or recolonization events. We also find evidence that the colonization of the Americas has resulted in introduction bottlenecks. However, while each sampling location shows evidence of a past population contraction and subsequent recovery, our results suggest that the bottlenecks in America have led to a reduction in genetic diversity of only ∼35% relative to African populations, and the American samples have retained high levels of genetic diversity (expected heterozygosity of ∼0.02 at synonymous sites) and have experienced only a minor reduction in the efficacy of selection. These results evoke the image of an invasive species that has expanded its range with remarkable genetic resilience in the face of strong eradication pressure.

## Introduction

Invasive species pose a threat to native species and ecosystems as well as human health and agriculture (Chornesky and Randall 2003; Paini et al. 2016; Dueñas et al. 2021). While many invasive species spread by displacing or outcompeting native species, many instead take advantage of under-utilized niches, often similar to those in their home environments (Elton 1958; Baker 1974; Sakai et al. 2003; Barrett 2015; Liu et al. 2020; Baker and Stebbins). One such pathway is developing an association with humans, either by exploiting anthropogenic changes to the ecosystem, or directly through the evolution of human preference (Hulme-Beaman et al. 2016). By adapting to these novel environments, invasive species face dramatic reductions in genetic diversity and the efficacy of selection, however they are nevertheless able to spread and outcompete native species, a long-studied conundrum dubbed the paradox of biological invasions (Sax and Brown 2000; Frankham 2005; Schrieber and Lachmuth 2017). For invasive species with human preferences, the shortcomings associated with the classic paradox of biological invasions may not hold true if they were adapted to human environments prior to their invasion, as the need to adapt to a novel environment may be greatly diminished or completely eliminated (Lee and Gelembiuk 2008; Hufbauer et al. 2012). The idea that some invasive species had previously adapted to humans and their environments is especially crucial for understanding the history and genomic consequences of invasive range expansions in human diseases and their vectors (Hufbauer et al. 2012; Powell 2019; Comeault et al. 2020).

Over the course of an invasion, populations of invasive species face dramatic demographic changes through their introduction and along their expansion front (Sakai et al. 2003). These include introduction bottlenecks that reduce genetic diversity to an extent that depends on the severity of the bottlenecks, the number of bottlenecks and the diversity of alleles they sample, and the amount of gene flow across the species range (Nei et al. 1975; Dlugosch and Parker 2008; Blackburn et al. 2016). The expected reduction in effective population size, and subsequent loss of genetic diversity following the introduction of an invasive species will impact the species’ ability to adapt (Barton and Partridge 2000; Brandvain and Wright 2016). During a bottleneck, the lower effective population size reduces the efficacy of selection relative to drift, allowing weakly deleterious mutations to drift to higher frequencies, and will also decrease the probability that beneficial mutations will fix in a population. Species facing anthropogenic selective pressures might experience such strong selection that adaptation can proceed in spite of reductions in genetic diversity; however, adaptation will still be slowed if the introduced population’s loss of diversity is severe enough that it will have to wait for new beneficial mutations (Orr and Unckless 2014; Osmond et al. 2019).

Importantly, the amount of gene flow between populations will also affect the efficacy of selection in contrasting ways: introducing new migrants to a population will increase the effective population size and thus the efficacy of selection, while the rate of influx of new mutations to a given locale can lead to a swamping effect, wherein selected alleles are overwhelmed by migration (Haldane 1956; Lenormand 2002; Yeaman 2022). For populations that are locally adapted, this gene flow is likely to primarily introduce maladapted alleles, but for populations with shared selective pressures like those from anthropogenic sources, gene flow may accelerate adaptation by introducing beneficial alleles that have arisen elsewhere in the global population that may not be present locally (Edelman and Mallet 2021; Yeaman 2022). In short, while theory predicts a decrease in the efficacy of selection and the rate of adaptation following a bottleneck and expansion, the degree to which this occurs in natural populations is less straightforward.

*Aedes aegypti*, the yellow fever mosquito, is a globally invasive species and the primary vector for the arboviruses dengue, yellow fever, chikungunya, and Zika. The species originated in Africa, where it spread during the African humid period, and differentiated into a generalist, *Ae. aegypti formosus* (hereafter *formosus*) and a human-adapted form, *Ae. aegypti aegypti* (hereafter *aegypti*) roughly 5,000 years ago (Brown et al. 2014; Rose et al. 2020; Rose et al. 2023). The *aegypti* form shows a higher preference for human hosts and an increased tolerance for rainfall variation, likely as a response to its strong association with humans (Rose et al. 2020). *aegypti* are relatively poor dispersers by their own flight, but their association with humans leads to long-distance dispersal, and high egg desiccation tolerance and dormancy means populations do not need to be immediately established, but may emerge years after their initial introduction (Machado-Allison and Craig 1972; Fischer et al. 2019; Mayilsamy 2019), possibly with diverse lineages from these egg banks (Kaj et al. 2001; Evans and Dennehy 2005). *aegypti* has recently spread worldwide, first into the Americas shortly after European colonization, and later into Asia (Brown et al. 2014). Worldwide *aegypti* populations appear to harbour far less variation than populations of *formosus* or mixed ancestry in Africa, and potentially result from a single origin of human preference (Gloria-Soria et al. 2016; Lozada-Chávez et al. 2023). The expansion of *aegypti* out of Africa was likely complex, with high ship volume to the Americas during the human slave trade offering ample opportunities for multiple early introductions and continued gene flow (Powell et al. 2018). In recent decades, increasingly globalized trade may have allowed for additional long-distance dispersal events.

In the Americas, *aegypti* has had a substantial impact on human health and history (Tapia-Conyer et al. 2009; San Martín et al. 2010; Shepard et al. 2011; Cafferata et al. 2013). Outbreaks of yellow fever were first reported in the 17th century (Blake 1968; Bryan et al. 2004), although they may have begun earlier (Carter 1931), and continued outbreaks in the following centuries wreaked havoc on indigenous populations and naïve Europeans. Various epidemics of dengue, beginning in the 17th century (Brathwaite Dick et al. 2012), chikungunya, beginning in the 19th century (Brathwaite Dick et al. 2012), and more recently Zika in the 21st century (Chang et al. 2016) have occurred and continue to occur throughout the Americas, including the United States. During the 20th century, several countries in South America began an eradication program for *aegypti*, imposing intense pesticide pressure on populations, and by 1962, 18 South American and Caribbean countries had reported eradication (American Health Organization 1997). Recolonization, recrudescence from unidentified populations, or a combination of both led to the reemergence of the vector across the continent (Kotsakiozi et al. 2017). Insecticide resistance evolved rapidly and repeatedly throughout the Americas, with several haplotypes underlying resistance to multiple insecticides reported in the literature (Kawada et al. 2014; Al Nazawi et al. 2017; Haddi et al. 2017; Saavedra-Rodriguez et al. 2018; Fan et al. 2020; Love et al. 2023).

Introductions into the United States have been more recent, perhaps earliest in the Southeast. The first confirmed collection of the species was done in Savannah (GA) in 1828 (specimen cataloged as *Culex taeniatus* Wiedemann; (Christophers 1960; Eisen and Moore 2013) but earlier outbreaks of diseases consistent with yellow fever in Spanish Florida in 1649 and the Northeast in 1668 suggest an earlier presence of the vector (Patterson 1992; Eisen and Moore 2013). These colonizations were then followed by subsequent spreads to the west, and multiple independent introductions reported in California with breeding populations first reported in the state in 2013 (Pless et al. 2017). Despite close monitoring of recent introductions and efforts to monitor and control populations throughout the Americas, previous genetic studies of the species have primarily focused on individual regions, high-level descriptions of population structure, and/or have used limited genomic information. Currently there lacks a detailed understanding of the demography of *aegypti* in North and South America, in particular the number and severity of *aegypti* introductions, the degree to which gene flow may aid in the spread and adaptation of populations throughout the continents, and how the efficacy of selection in introduced populations of *aegypti* has been affected by their demographic history.

Here we examine a set of 131 whole genomes from African and American *Ae. Aegypti* populations to model the demographic history of the species’ spread into the Americas, and investigate its effects on genome-wide diversity and the efficacy of selection. First we investigate patterns of population structure across our dataset, within and among Africa and the Americas. We then infer the history of effective population size changes and split times between populations within and between the Americas and Africa. Finally, we investigate the efficacy of selection of all populations by examining levels of diversity genome-wide and in protein-coding genes and by inferring the distribution of fitness effects of new mutations. We find that the spread of *Ae. aegypti aegypti* to the Americas appears to be best characterized by multiple apparent introductions and limited subsequent gene flow at fine and coarse geographic scales, and that the history of bottlenecks and strong selection in the introduced range has had genome-wide impacts on diversity and the efficacy of selection. Despite a 33-40% reduction in neutral genetic diversity, *aegypti* in the Americas maintains high diversity and appears to have experienced only a modest reduction in the efficacy of selection. The limited effect of its introduction history and recent anthropogenic selection on diversity and the efficacy of selection illustrates a surprisingly resilient species at the genomic level—one that poses a threat to future eradication efforts. We discuss the implications of this history and its genomic impact in the context of invasive species and vector control, and adaptation (past, ongoing, and future) in the species.

## Results and Discussion

We used publicly available sequencing data from several studies, recently collated in (Love et al. 2023). This dataset includes 27 specimens from California and Florida (Lee et al. 2019), 18 from Santarem, Brazil, 13 from Franceville, Gabon, 19 from Kaya Bomu, Kenya, and 20 from Ngoye, Senegal (Rose et al. 2020), and 10 from Cali and 24 from Río Claro, Colombia (Love et al. 2023), for a total of 131 specimens, 79 of which are from the Americas. Hereafter we refer to specimens sampled from the same location as accessions, and to groups inferred to show evidence of recent shared ancestry as populations or clusters, depending on context. The African accessions used here were previously scored for human preference, with the Gabon accession exhibiting little-to-no human preference, the Kenya accession exhibiting intermediate/mixed preference, and the Senegal accession exhibiting strong human preferences (Rose et al. 2020), while the American accessions are assumed to exhibit strong human preferences.

### *Aedes aegypti* is highly structured at fine and coarse scales

To characterize the population structure of our samples, we first used principal component analysis, beginning with our full dataset. The first and second principal components, representing 12.35% and 7.25% of the variance, respectively, separate the African accessions from the Americas and from each other (Figure 1B; scree plot in Supplementary Figure 1). The African accessions separate distinctly from each other, with more apparent diversity in Kenya than in either Gabon or Senegal. The American samples cluster together, with the three South American accessions and the USA accessions clearly differentiated within this regional cluster. Among the African samples, Senegal appears to be most closely related to the American samples, and particularly to samples from the US, consistent with previous studies identifying Senegal as a proxy for the ancestral *aegypti* form, and with a more recent invasion in the US. However, one Senegal specimen appears to cluster near to the Gabon cluster, which has been previously noted (Rose et al. 2020). This specimen could represent a recent migrant from a population closely related to the Gabon accession, or a generalist individual within a larger-than-recognized range of *formosus*. PC1 is consistent with separating accessions by either latitude or host preference, and interestingly, PC2 primarily separates Kenya from Gabon and Senegal. Kenya and Gabon both exhibit primarily low human preference, with the former composed of mixed preference, evident here with the spread of the Kenyan sample along both PCs, with some specimens closer in PC-space to American samples.

**Figure 1:**
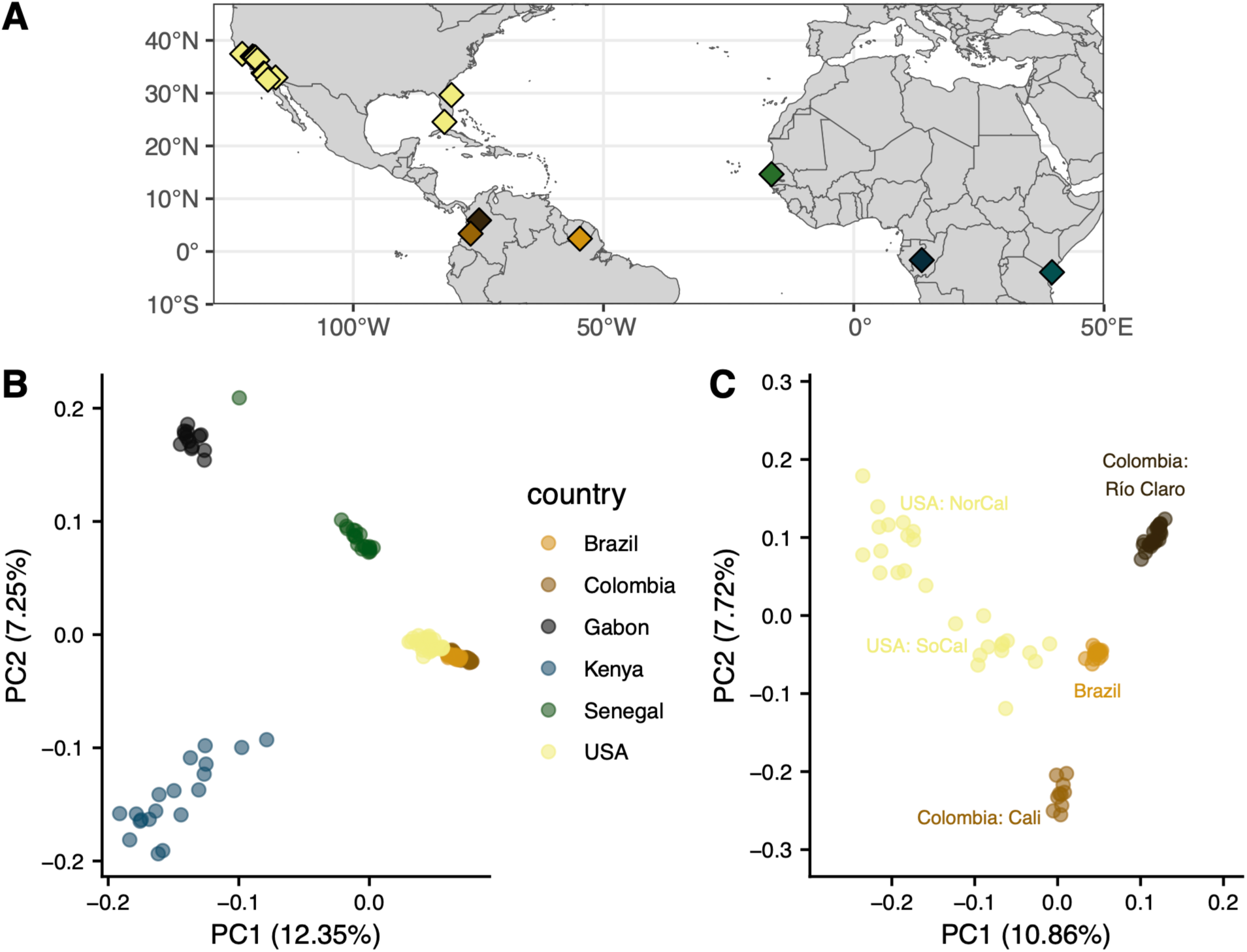
(A) Sampling locations of specimens used in this study. (B) Principal component analysis (PCA) of all specimens highlighted by country of origin. (C) PCA of American samples highlighted by location.

Within the Americas, principal components reveal more complex structure (Figure 1C; scree plot in Supplementary Figure 2). Accessions from Colombia and Brazil separate clearly, with Brazil closer to either Colombian accession than the Colombian accessions are to each other, as previously reported (Love et al. 2023), despite being relatively close in geographical space (Figure 1A); indeed, despite being separated by only ∼500km, these two accessions are found nearly on opposite ends of PC2. As may be expected from geographically spread samples with few from a single location, the samples from the US are loosely clustered into two groups in the first two PCs, and three to four groups apparent in deeper PCs (Supplementary Figure 3), similar to what was previously reported with these samples (Lee et al. 2019). The first US cluster in PC 1 and 2 groups Northern California and some accessions from Central California in the upper left of Figure 1C. The second cluster, closer to Brazil in the centre of Figure 1C, consists of Southern California, a subset of accessions from Central California, and Florida. In PCs 3 to 5, Central and Northern California clearly separate into groups: a cluster of Menlo Park with Madera and Fresno, and a cluster of Clovis and Sanger (Supplementary Figure 3). This separation is remarkable in that Clovis and Sanger are roughly 250 km apart, and on either side of Fresno with which they do not cluster. The Southern California and Florida cluster is largely maintained at PCs 3-5, with only the Florida sample from Vero Beach separating beginning at PC 4. The accessions from Exeter, California, located in the Central Valley south of Fresno, cluster closely with an accession from Key West, Florida, and near to the Southern California accessions. Similar to the Northern California cluster, structuring within the primarily Southern California cluster is hallmarked by a combination of fine and coarse scale geographic separation, and can likely be clustered into two to three groups within California, and two separate Florida accessions.

To more explicitly model population structure within our dataset, we used ADMIXTURE. ADMIXTURE clusters samples using allele frequencies into a pre-defined number of groups, modeling each individual as a mixture of the clusters. Using the whole sample set, the best number of clusters according to cross validation is *K*=2, where samples are largely split into African and American groups (Figure 2A). Several specimens in Kenya and nearly all in Senegal exhibit substantial admixture proportions, with the major component grouping with Gabon and the minor component with the Americas. All specimens from Gabon are entirely assigned to the same group (the major ancestry component in Africa), while the Brazilian and both Colombian accessions are all completely assigned to the opposing group—the minor ancestry component of Kenya and Senegal. Specimens from the US are nearly entirely composed of ancestry shared with South America and the minor ancestry component of Kenya and Senegal, though three specimens, one of each from Southern California, Central California, and Vero Beach, Florida, show small amounts of the Gabon-like ancestry.

**Figure 2:**
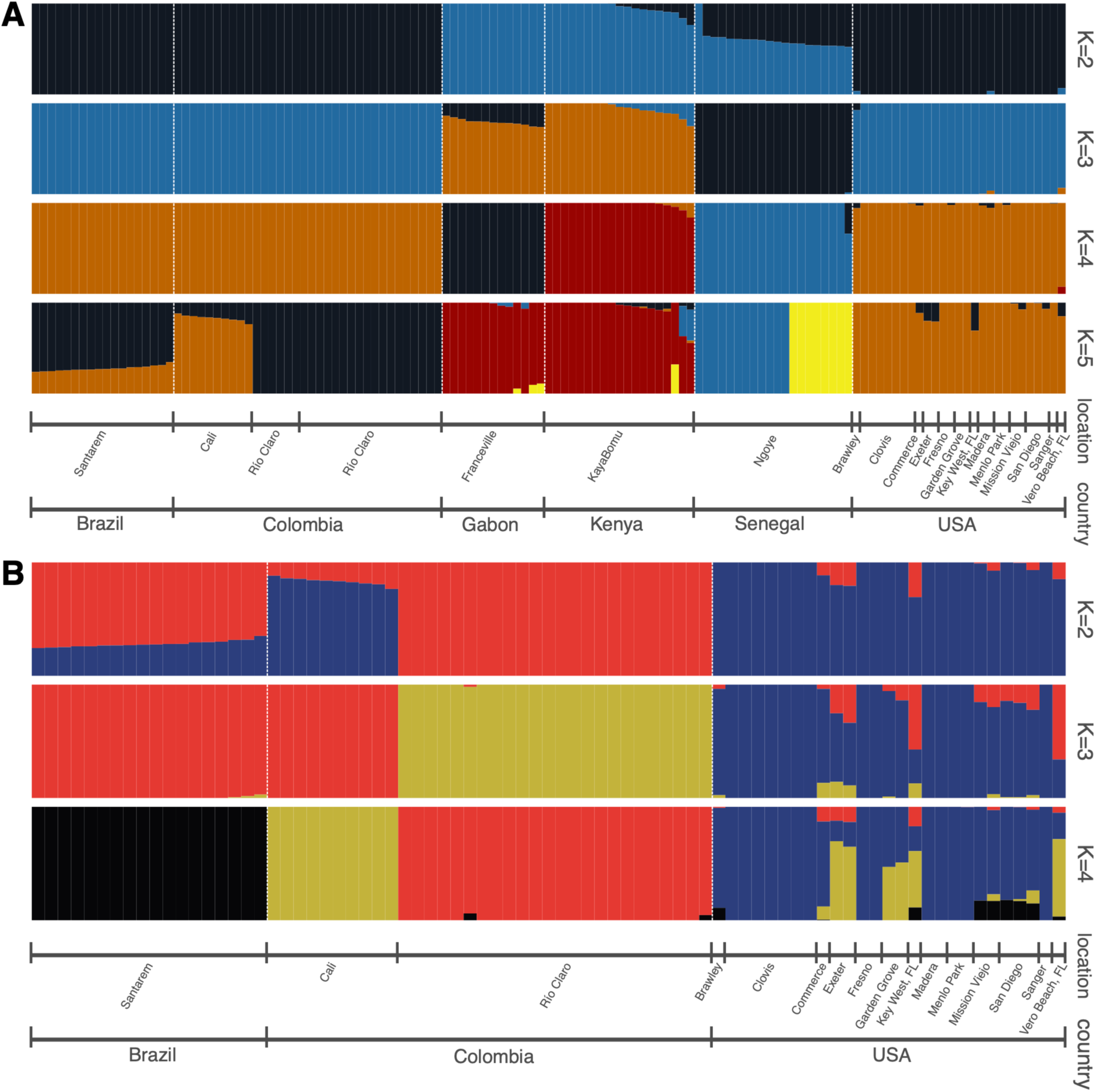
Admixture results for global samples (A) and for the Americas (B). The best *K*, obtained via cross validation, is *K*=2 for both (A) and (B), but we show results for higher *K* for additional context.

Mixed ADMIXTURE group membership in Kenya and Senegal is consistent with domestic ancestry or human preference previously reported in these accessions. Senegal, apart from one unadmixed specimen assigned entirely to the Gabon cluster, varies between roughly 50:50 and 70:30 split of Gabon-like to South American-like group memberships, while about half of Kenyan samples have a small amount of South American-like ancestry. Higher values of *K*, although a slightly poorer fit according to cross validation (Supplementary Table 1), provide some context for the minor ancestry components in Senegal and Kenya. At *K*=3 and above (Figure 2A), the minor ancestry in Kenya groups with the American accessions, while Senegal largely appears unadmixed and groups with some ancestry in Gabon, or with itself, until K=5 where it splits into two groups shared in minor proportions with some specimens in Gabon and Kenya. This is concordant with Senegal grouping most closely to the American accessions in PCA, but some Kenyan samples being nearly equidistant. While the extant accession in Senegal may represent the nearest known representative of an ancestral population to the American samples, these results from PCA and ADMIXTURE are consistent with ancestral population structure resulting ancestry shared among some African populations, including some domestic *aegypti* haplotypes that spread to the Americas (Slatkin and Pollack 2008). Back migration from the Americas to African populations (Brown et al. 2011; Powell and Tabachnick 2013) has also been proposed and would be consistent with low levels of American-like ancestry in Kenya during a second wave of the shift to human preference in African cities (Rose et al. 2020).

Within the Americas, the best number of groups according to cross validation is again *K*=2 (Figure 2B), and the patterns of admixture are identical to the patterns in the Americas when considering all samples at *K*=5 (Figure 2A). At *K*=2, accessions from Brazil and from Cali, Colombia are shown as admixed, with reciprocal minor ancestry components, while Río Claro, Colombia is unadmixed of the major ancestry component in Brazil. Greater grouping between Brazil and Río Claro is consistent with results from PCA (Figure 1C) and divergence statistics (Supplementary Figures 4 and 5 for genome-wide *F_ST_* and *D_xy_*) showing greater divergence within Colombia than between Río Claro and Brazil. The US is represented as partially admixed and primarily composed of the major ancestry component in Cali. The admixture in the US is isolated to the previously mentioned cluster present in the first two PCs of Southern California, Florida, and the Central California accession of Exeter (Figure 1C), and is present in some but not all of these individuals at *K*=2, varying from minor to sizable proportions. At higher values of *K*, population structure remains qualitatively similar, with additional groupings adding to the diversity of admixture within the admixed US specimens, while the remaining US specimens compose their own group and South American accessions become unadmixed and quickly split among each other. This coarse-scale separation again reflects the two major clusters in the Americas, with little effect of geographic distance as evident in the separation within Colombia and the mixed fine and coarse-scale structure in the US.

In summary, both ADMIXTURE and PCA results from the Americas depart strongly from expectations under a simple model of geographic isolation by distance. This may lend support to the possibility that there have been several independent introductions to the Americas, each sampling a subset of the ancestral *aegypti* variation available in Africa. We examine this possibility more directly in the section below.

### American *aegypti* underwent multiple, strong bottlenecks

We sought to investigate the number and timing of introductions of *Ae. aegypti* among our sampled accessions, which represent a range of events spanning from among the earliest to the most recent proposed introductions to the Americas. We began by inferring the historical effective population sizes among all countries using SMC++, making use of recent estimates for the generation time and mutation rate of *Ae. aegypti* (Rose et al. 2023). We report results obtained using the same model regularization parameter (rp=5), which provided the most consistent fit across the seven groups examined in Figure 3 as evidenced by coalescent simulations (Supplementary Figures 8–14). These analyses suggest that the ancestral effective population size (*N_e_*) among all accessions must have been between 300,000 and 400,000 until about 20,000 years (300,000 generations) ago (Figure 3A & B). All accessions experienced a reduction in *N_e_* around 10,000 years (150,000 generations) ago, coinciding with the end of the African humid period as previously reported (Rose et al. 2023). Kenya and Gabon recovered to near ancestral *N_e_* over a span of a few thousand years, while Senegal appears to have suffered a deeper bottleneck than the other African accessions, dropping to roughly 80,000, with the lowest *N_e_* point around 1,000 years (15,000 generations) ago, before recovering to about 200,000. This bottleneck in Senegal occurred between 5,000 and 10,000 years ago, consistent with the founding and diversification of the *aegypti* domestic ancestry within Africa in the dry, variable Sahel after the African humid period (Rose et al. 2023). As an alternative approach to estimating population size histories, we also ran Stairway Plot 2 (Methods), and obtained qualitatively similar patterns as those obtained via SMC++ in terms of the differences relative severity of bottlenecks across accessions (Supplementary Figure 6).

**Figure 3:**
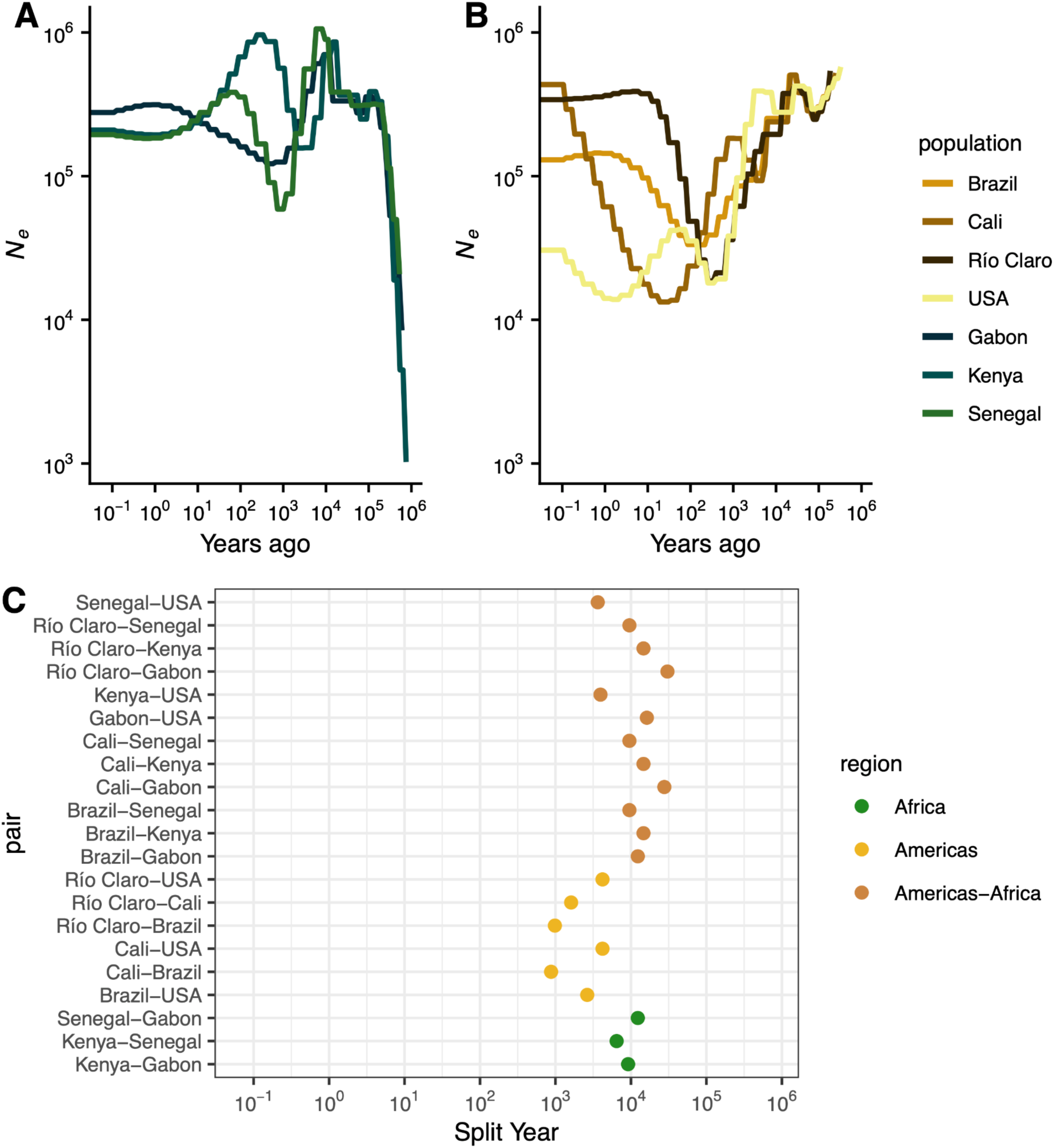
Estimated population size (*N_e_*) history and split times from SMC++. (A) *N_e_* histories for all African accessions. (B) *N_e_* histories for all American accessions. (C) Split times for all accession pairs.

Among the American accessions, we infer strong, prolonged bottlenecks that vary in their timing and intensity. All American accessions began to decline between 5,000 and 10,000 years ago, and continued to decline for thousands of years (Figure 3B). In the accession from Cali, Colombia, we infer a fluctuating *N_e_* that was approximately stable until about 700 years ago, when its introduction bottleneck started and continued to decline until it reached a low *N_e_* of 13,312 only 20 years (300 generations) ago, followed by a recovery. The relatively larger Ne of the Cali accession that is inferred at the time when other American accessions began to exhibit a contraction may suggest a more diverse sampling of lineages, possibly through multiple introductions or migration after colonization; both of these possibilities would result in a period of decelerated coalescence and are consistent with the higher degree of admixture inferred in Figure 2B for this accession than the other South American accessions. In Río Claro and Brazil, bottlenecks beginning after the Africa humid period continued until their lowest *N_e_* points of 18,639 and 33,225 occurred 250 and 150 years (3,750 and 2,250 generations) ago, respectively. In all three accessions we infer massive recoveries, to a present day *N_e_* of 433,679, 340,611, and 129,800 in Cali, Río Claro, and Brazil, respectively. Methods based on the sequentially Markovian coalescent like SMC++ have limited ability to accurately estimate the *N_e_* in very recent time, however the qualitative result of rapid recoveries in these accessions occurs within the span of generations in the past where these methods have reasonable accuracy (Patton et al. 2019), at least in Río Claro and Brazil, and is consistent with monitoring and disease prevalence suggesting a rebound after the collapse of Pan-American eradication efforts (reviewed in (Webb 2016)), potentially through recolonization (Monteiro et al. 2014). While the bottlenecks in each locality are inferred to have occurred at different times over a span of roughly 200 years, we cannot be confident that they all represent different events; however, we can be confident that each of the South American accessions experienced strong bottlenecks and recoveries in the very recent past. Introduction bottlenecks in the South American accessions resulted in reductions in *N_e_* of 94%, 89%, and 96% in Río Claro, Brazil, and Cali, respectively, which appear to have occurred no later than 4,000 generations or 250 years ago.

As evidenced by the PCA and ADMIXTURE results, the United States accessions appear to represent a complex structuring of populations. However, grouping the US accessions into a single population when running SMC++ results in reductions in inferred *N_e_* similar to those in Cali, pointing to substantial bottlenecks in the underlying structured populations. We therefore split the US into five approximate populations representing Southern California, Northern California, Florida, Clovis and Sanger, CA, and Exeter, CA, following clustering in PCA space (Supplementary Figure 3) and results from previous studies (Lee et al. 2019). All populations experienced a substantial bottleneck in the recent past, all beginning roughly 10,000 years ago as in the South American populations, but all peaking at a similar *N_e_* of about 5,000 only 55 years (825 generations), except for Florida which peaked about 240 years ago at about 15,000 (Figure 4A). All populations similarly are inferred to have recovered to between 400,000 and 1,500,000 at the present day, except for Florida and Exeter which are inferred to have plateaued 5 to 10 years ago around 100,000, although this lower recovery may be an artifact of having only two samples in each of these populations. Similarly, the samples from Florida represent divergent lineages (Supplementary Figure 3), which may inflate *N_e_* estimates and bias the timing of bottlenecks. Among California clusters, the timing of the start and peak of each bottleneck is similar, though the Northern California cluster is slightly shifted to the more recent past, consistent with previous reports of a later, independent introduction relative to Southern California (Lee et al. 2019). As a whole, US populations experienced deeper and more recent bottlenecks than the South American populations, consistent with later introductions, and an earlier introduction in Florida than in California, though the differences in *N_e_* between the US clusters are minimal. The apparent earlier introduction in Florida with a lowest *N_e_* roughly 240 years ago is after historical records of diseases matching the symptoms of Yellow Fever first appeared in Spanish Florida and the Northeast (Patterson 1992; Eisen and Moore 2013), but shortly before reports of the species in Georgia (Christophers 1960; Eisen and Moore 2013) and dengue outbreaks throughout port cities in the Eastern US (Brathwaite Dick et al. 2012).

**Figure 4:**
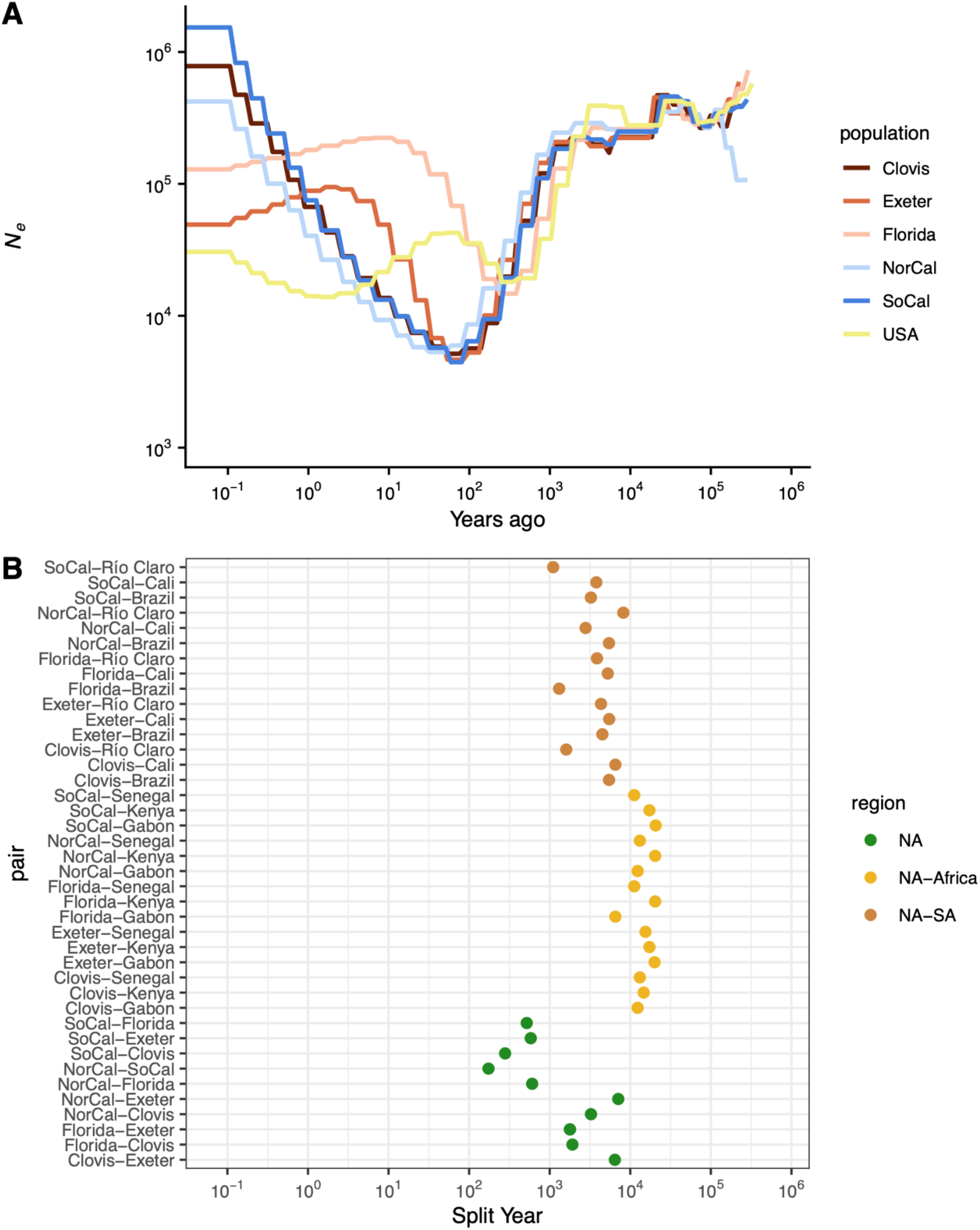
(A) Effective population size histories for approximate USA populations, with the estimated history of the combined accessions from the previous plot. (B) Split times for all approximate USA populations with all other accessions.

We also used SMC++ to infer population splits between all pairwise accessions. This model infers a split time using the within- and cross-population coalescent rates assuming a clean split model. At a broad scale, we find that split times involving African samples are older than split times within the Americas, though split times between Africa and the Americas span a range of time older and more recent than splits within Africa (Figure 3C). Among the African accessions, Senegal and Kenya are inferred to be most closely related, followed by Kenya and Gabon, and Senegal and Gabon. This order of population splits is consistent with previously published behavioural results (Rose et al. 2020), and support the notion that the Senegalese and a portion of the Kenyan accessions appear to represent a derived domestic *aegypti* ancestry, while the Gabon accession have maintained the ancestral generalist *formosus* form. The split times among the African accessions span a few thousand years, from 6,500 years ago for Kenya-Senegal, to 9,175 for Kenya-Gabon, and 12,300 years ago for Senegal-Gabon. Assuming a single origin for human preference, mixed *aegypti* and *formosus* ancestry in the Kenyan accession would likely be the result of recent migration from an *aegypti* population more closely related to Senegal. In a clean split model not accounting for this recent migration, the estimated split time would be closer to the present, rather than correctly inferring an older split followed by recent migration. Alternatively, as previously mentioned, the domestic ancestry in these accessions may have sampled different diverse haplotypes in the ancestral domestic population, yielding the same signal, consistent with ADMIXTURE results.

Between Africa and the Americas, we infer a broad range of population split times. Among the South American accessions, we infer nearly identical split times in increasing order to Senegal (split times between the three South American accessions and Senegal were each estimated to be 9,545 years ago), Kenya (split times all estimated to be 14,678 years ago), and Gabon (27,699 years to Cali and 30,479 years to Río Claro), except for the Brazil-Gabon split which is estimated to be slightly more recent than the Brazil-Kenya split (12,343 years ago; Figure 3B). These split times far predate their introduction to the Americas, and are on the order of or older than the splits within Africa, suggesting a deep split between extant African populations and the ancestral domestic population that eventually invaded the Americas. In contrast, split times between the US and Africa are much more recent, though still far predating the American introduction (3,636 years ago with Senegal, 3,953 years ago with Kenya, and 16,256 years ago with Gabon); aside from a more recent split, this result could also be explained by migration between the African populations and the lineages leading to the US accessions after the split between the South American and African samples.

Within the Americas, estimated split times vary by thousands of years. In South America, our ordering of estimated population splits is consistent with results from PCA: Cali and Brazil most recently split (875 years ago), followed by Río Claro and Brazil (983 years ago), and finally Río Claro and Cali (1,613 years ago). These splits are well before their introductions to the Americas, and differences among them will reflect differential inheritance of ancestral variation and any migration after their introduction, but strongly suggest independent introductions to each location. Split times between South America and the US are deeper: estimated splits between the US and both Río Claro and Cali are identical (4,203 years ago), while the split between the US and Brazil is more recent (2,636 years ago). Again, the differences between these split times may reflect ancestral sorting, and the extent of recent migration (if any). However, in this case our estimates may also be affected by the presence of population structure in the US.

With the potential impact of population structure in mind, we also considered split times for five different clusters of the US accessions to all other accessions (Figure 4B). When considering these US partitions separately, the estimated split times are much deeper than when considering the US as a whole. The splits between US clusters and African accessions are slightly older than those estimated between South American and African accessions. However, the split between Florida and Gabon is estimated to be substantially more recent at 6,536 years ago, potentially related to the more recent splits between Brazil and Gabon (Figure 3B) and Florida and Brazil (Figure 4B). For most US clusters compared to Cali and Brazil, the estimated split times are 4,500 to 6,500 years ago; however, Florida, Southern California, and Northern California have more recent split estimates, driving the US splits to these accessions as a whole towards the present. Southern and Northern California both have estimated splits to Cali about half as long ago as the rest of the US (3,791 and 2,790 years respectively), and Southern California also has a more recent estimate to Brazil (3,250 years ago). Florida stands out with a much more recent split estimate to Brazil (1,310 years ago), more recent even than its split times to Exeter and Clovis. While these estimates are much older than reasonable introduction times to the Americas, they may reflect a similar ancestral source population, or an unsampled South American source population for Florida related to the accession presented here, especially in concordance with the high levels of admixture observed in the Floridian samples (Figure 2B). In total, the estimated split times between the US partitions and those in South America do not suggest an origin for the sampled US accessions in related South American populations.

Within the US, estimated split times again suggest a complex introduction history (Figure 4B). There is a strong stratification in split times between a few US clusters—Florida and Northern California both have deep split times to the Central California accessions in Clovis/Sanger and Exeter, similar to the deep split between Clovis/Sanger and Exeter. These deeper splits span a broad range of time (1,785 to 7,100 years ago) with Florida more closely related to Clovis/Sanger and Exeter, while Northern California and Exeter is the deepest split. While deep splits between Florida and Central California may not be surprising, and are consistent with independent introductions to California and Florida, the deep splits between some of the more geographically close California samples are more unexpected. For example, the accessions of Clovis/Sanger and that of Exeter are separated by about 80km (split time estimate: 6,444 years ago), and Clovis/Sanger and Fresno (part of the Northern California cluster), are only separated by 16km (split time estimate: 3,255 years ago). These findings suggest that there has been long distance dispersal of related haplotypes within California, along with multiple introductions of lineages to several cities in the state that appear to have experienced little-to-no gene flow since their introduction and up until the time of sampling.

Among the more closely-related set of accessions in the US, estimated split times leave their origins less clear. The most recent split is between Northern and Southern California at 174 years ago, suggesting either an introduction from elsewhere within the Americas for both sets of accessions, as has been suggested at least for the Southern California populations (Lee et al. 2019), or a split somewhere within the Americas and subsequent dispersal to different regions. Southern California and the Clovis accession are estimated to have split similarly recently at 280 years ago, again suggesting they may have originated through a split from a population already within the Americas, or possibly long-distance migration along the California Highway 99 corridor. The remaining splits, between both Northern and Southern California and Florida, and Southern California and Exeter occur within a timespan ranging from 519 to 606 years ago. While these timeframes are older than a likely introduction, they remain much more recent than splits within South America or between American and African accessions, suggesting divergence within the Americas is not unlikely, but also consistent with separate introductions followed by gene flow from the same source population.

### Introduced populations suffer minimal reductions in selection efficacy despite substantially reduced diversity

Given the strong bottlenecks experienced by all sampled introduced accessions, we sought to quantify the impact of the introduction history and previously described history of recent strong selection (Love et al. 2023) on genome-wide levels of diversity. We first estimated genome-wide neutral diversity using 4-fold degenerate sites (*π*_4_) (Figure 5A). Among African accessions, neutral diversity ranged from about 0.03 in Senegal to nearly 0.035 in Kenya. This level of diversity is similar to that reported in *Anopheles gambiae* (Corbett-Detig et al. 2015) and somewhat higher than previous genomic estimates of all sites (Rose et al. 2020; Love et al. 2023). As expected given the demographic history of the accessions, neutral diversity in the introduced range was much lower than in the ancestral range: nearly 0.02 in Brazil and Cali, Colombia, and ∼0.018 in Río Claro, Colombia. Diversity in the combined USA set of accessions was still less than 0.025, despite being inflated by population structure. While a 33-40% reduction in diversity relative to Senegal is substantial, it is perhaps less of a reduction than expected given the large bottlenecks and eradication efforts sustained by these accessions. In an absolute sense and relative to other species (Leffler et al. 2012; Buffalo 2021), neutral diversity near 0.02 in the Americas is high, posing potential problems for future control efforts.

**Figure 5:**
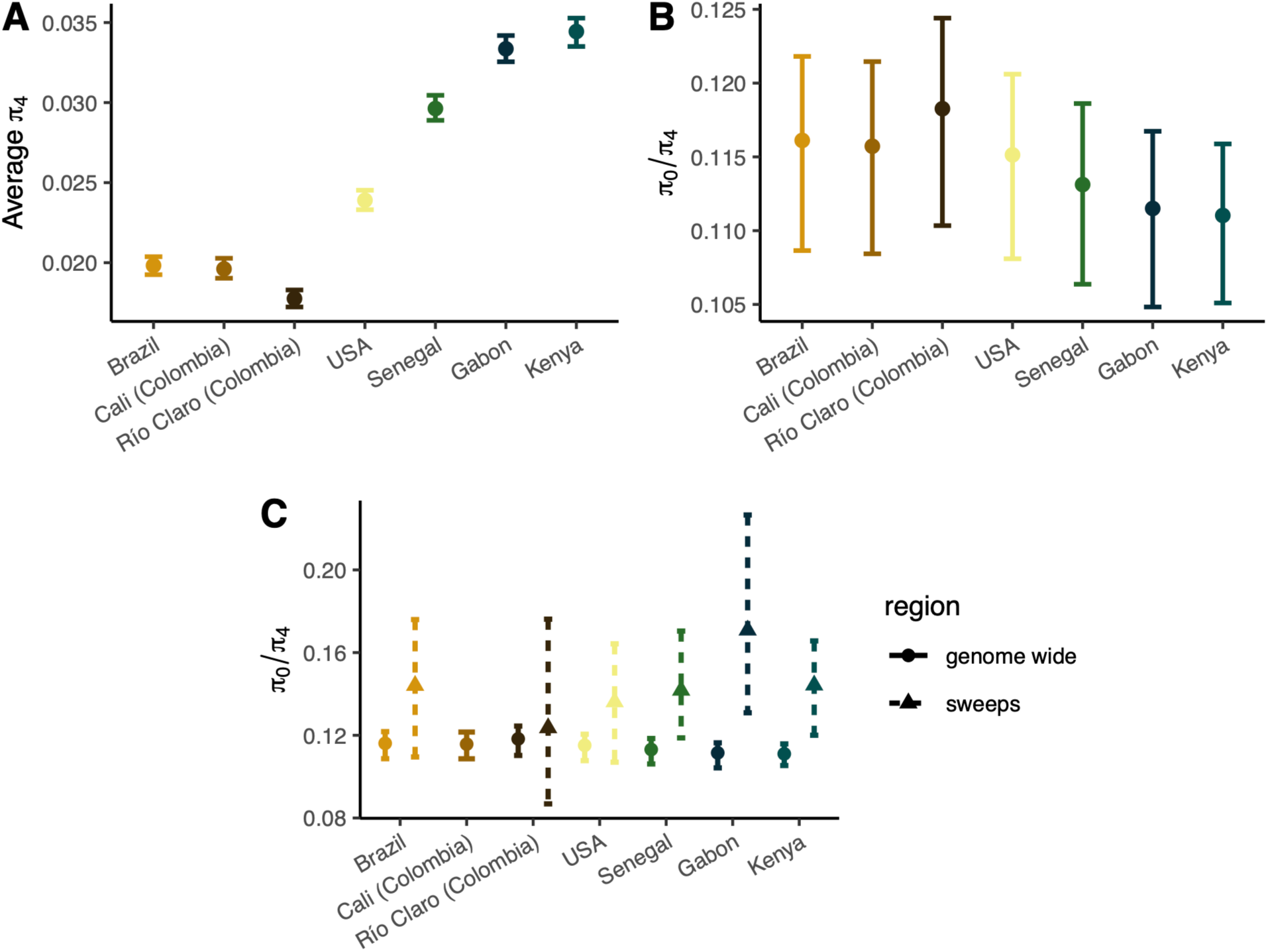
Diversity and measures of the efficacy of selection. (A) neutral diversity, *π*_4_. (B) *π*_0_/*π*_4_ with bootstrapped errors for all accessions. American accessions show a subtle decrease in the efficacy of selection. (C) *π*_0_/*π*_4_ in regions around sweeps (note that Colombia was combined for sweep scans, so is reported as a combined value for Río Claro and Cali, and as such, estimated neutral diversity is likely overestimated in the sweep regions, yielding an artificially lower *π*_0_/*π*_4_).

The combined effect of the introduction bottleneck(s) and recent selective pressures in the American *Ae. aegypti* populations should feed back into its ability to adapt because both of these phenomena are expected to reduce the efficacy of selection. We sought to test this using two indirect measures of the efficacy of selection. First, we estimated the ratio of 0-fold to 4-fold diversity in each accession (Figure 5B). Assuming negative selection against new deleterious mutations is persistent, an increase in the *π*_0_/*π*_4_ ratio reflects a reduction in the efficacy of selection, i.e. an increase in the proportion of effectively neutral mutations as seen by selection has allowed weakly deleterious mutations to accumulate. We find African accessions to have a relatively low *π*_0_/*π*_4_ ratio (∼0.111 in Kenya and Gabon, and ∼0.113 in Senegal). Consistent with a reduction in the efficacy of selection, the introduced accessions have higher ratios, near 0.116 in Brazil and Cali, and 0.118 in Río Claro, while the grouped USA accessions were near 0.115 (here artificially deflated due to population structure). These values are consistent with those reported in other insect species with no known history of introductions or eradication efforts (Chen et al. 2017). This decrease in the efficacy of selection is subtle, though perhaps unexpected given that neutral diversity in the introduced range remains high in an absolute sense, and in a relative sense was not as reduced as much as may have been expected.

Inferring changes to the efficacy of selection via *π*_0_/*π*_4_ in nonequilibrium populations can be complicated by differences in the recovery time to equilibrium between nonsynonymous and synonymous alleles (Brandvain and Wright 2016). As such, an increase in *π*_0_/*π*_4_ of effectively neutral mutations as we find here may not solely reflect a reduced efficacy of selection. To address this, we can also use explicit modeling of the distribution of fitness effects (DFE) of new mutations. When we infer the DFE in each accession, we find clear differences between the African and American accessions—American accessions, and to a lesser extent Senegal, exhibit both an increase in strongly selected mutations and nearly neutral mutations (Figure 6). The simultaneous shift to low and high Nes categories is evident in all introduced accessions, including the grouped USA accessions where the effect is attenuated by population structure (Andersson et al. 2023). The shift is primarily driven by a reduction in the low and intermediate effect classes, with minor increases in the nearly neutral class and large increases in the strongly selected class. In the nearly neutral category, we see increases in the American accessions of 1.7-3% over Senegal, and 3.5-4.5% over Kenya, representing a small but significant decrease in the efficacy of natural selection in introduced populations. Consequently, we do observe a small decrease in the efficacy of selection consistent with the increase in the ratio, though the biggest difference in the introduced accessions is in the unexpected shift towards more strongly deleterious mutations. However, it is possible that we have limited power to differentiate between moderately (10-100 *N_e_*s) and strongly deleterious mutations (100+ *N_e_*s) without knowledge of the true underlying DFE (Kousathanas and Keightley 2013); grouping these categories yields increases in the strongly selected categories for the American accessions of only 1-2% over the African accessions (Supplementary Figure 7).

**Figure 6:**
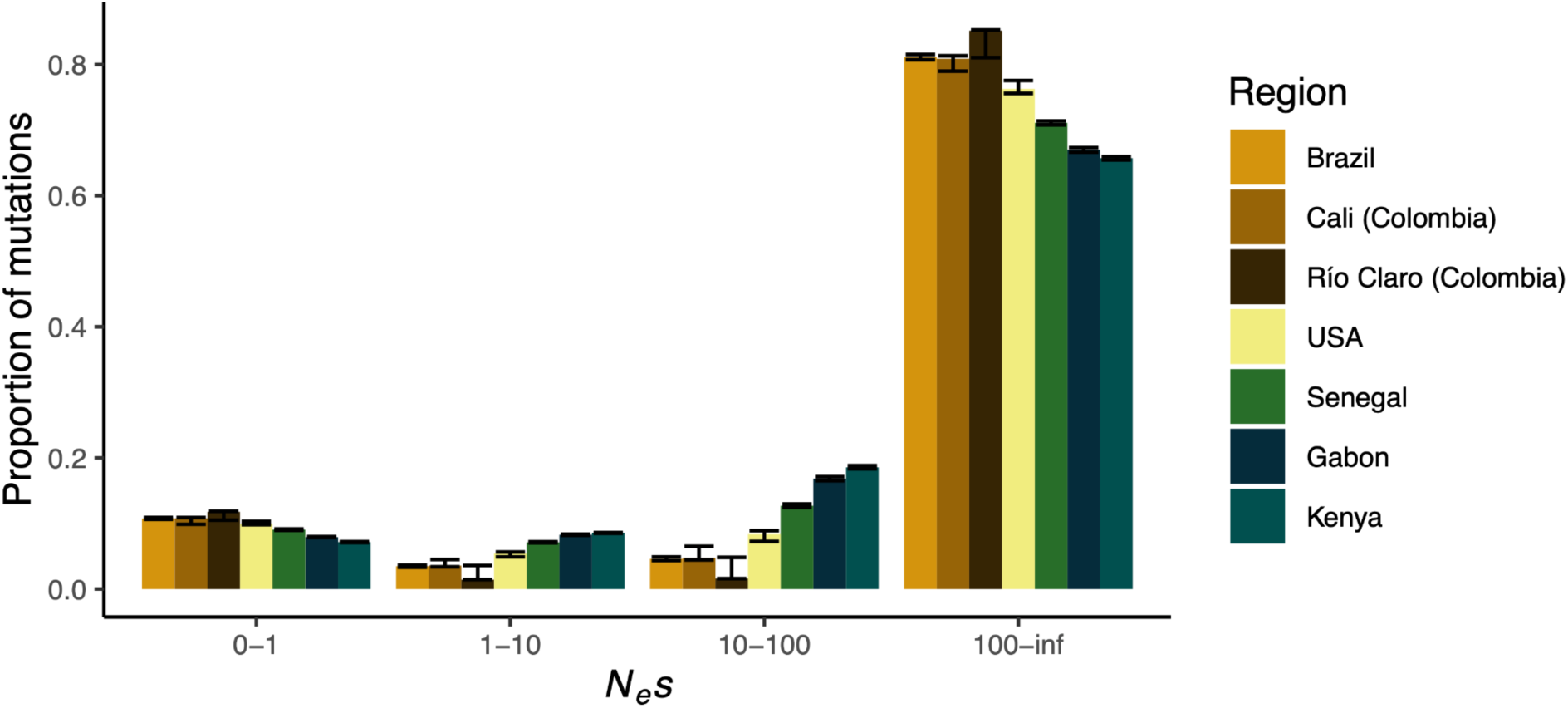
Distribution of fitness effects (DFE) of new mutations for 0-fold degenerate sites relative to 4-fold degenerate sites for all regions. Error bars represent 95% bootstrapped confidence intervals for each bin.

The efficacy of selection is additionally affected by selective sweeps (Smith and Haigh 1974), and in *Ae. aegypti*, particularly at sites under strong selection such as those underlying insecticide resistance (Love et al. 2023). While demography and selection both contribute to reductions in *N_e_*, selective sweeps only affect diversity in the regions linked to selected sites (Smith and Haigh 1974), meaning that the impact on both diversity and the efficacy of selection are heterogeneous along the genome. To elucidate the contribution of selective sweeps to the reduction in the efficacy of selection genome-wide, we estimated *π*_0_/*π*_4_ in the 10kb surrounding the top 1% of previously identified sweep signals (Love et al. 2023). We find the *π*_0_/*π*_4_ ratio is substantially increased in regions surrounding sweeps in all accessions (Figure 5C). The Colombian accessions do not show as much of an increase as in other accessions, though this is likely due to the Colombian accessions having been grouped together in the previous work, dampening the effects of independent sweeps. This result is consistent with a substantial draft effect in the species, though here we are unable to quantify the amount of the genome affected. Future work is needed to determine the relative contribution of demographic and selective processes in shaping genome-wide diversity and the efficacy of selection, with important implications for *Ae. aegypti*’s evolved response to control efforts.

### Conclusions

Here we investigated the demographic history of *Aedes aegypti* with respect to its introduction from Africa to the Americas. Our results suggest extensive population structure at fine and coarse scales, with some limited evidence for admixture in distant accessions in South America and between South America and some accessions in the US, while many accessions in close proximity exhibit deep divergence with no evidence of recent admixture. In particular, accessions from Colombia and Brazil appear to be largely genetically distinct, with the accession in Cali, Colombia more closely related to the accession from Santarem, Brazil than to the accession in Río Claro, Colombia, despite the relatively short geographic distance between the Colombian sampling locations. Similarly, accessions in California overlap each other in geographic space in the Central Valley, despite having estimated divergence times hundreds of years ago. The geographic overlap near Fresno, California separates the broad Northern California genetic cluster from the Southern California cluster, the latter of which includes accessions from Florida, further highlighting the complexities of scale in population structure of the species. Together, our observed patterns of population structure and admixture imply that gene flow between regions may be limited, but occasional new (or re-) colonizations likely occur through long distance dispersal, as is likely within California. Our admixture results are more broadly consistent with the sorting of diverse ancestries during the introduction bottlenecks, supported by deep estimated divergence times. In the context of the spread of beneficial mutations, such as those conferring insecticide resistance, our results suggest that adaptation may occur through parallel mutation instead of through the spread of single-origin mutations across large geographic ranges, though future work is needed to more directly test this hypothesis at loci of interest.

Regarding the number and timing of introductions in the Americas, we find through inferences of historical effective population sizes and population split times that populations in both South and North America likely originated through multiple introductions. We note that inferring the precise timing of introduction events with genomic data is challenging, and our estimates should not be treated as exact, however, our divergence estimates suggest American populations split thousands of years ago, long before the proposed introduction of *aegypti* to the Americas via the slave trade (Rose et al. 2023). Similarly, while the differences in bottleneck strength and timing, the general lack of admixture in the Americas, and estimated split times predating possible introduction times by hundreds of years strongly suggest there have been multiple origins in South America, we cannot directly ascertain whether populations that were declared eradicated were in fact eliminated and recolonized from elsewhere, or if instead recovery occurred from the expansion of a smaller number of individuals surviving in refugia. However, the fact that we do not observe especially strong bottlenecks that would be expected for a population recovering from near eradication (discussed below), perhaps makes the recolonization hypothesis more likely. Similarly, for accessions in the US, estimated split times and bottleneck severities and timings point to independent origins relative to the sampled South American accessions, although we cannot identify the introduction route. Notably, some pairs of North and South American clusters exhibit markedly more recent estimated split times than other such pairs (e.g. Southern California and Clovis relative to Río Claro, and Florida relative to Brazil). These more recent split time estimates, which we obtained under a model that does not allow for migration, could potentially be underestimates caused by post-split gene flow between some North and South American clusters; this notion is consistent with our evidence of relatively low but nonzero admixture proportions shown in Figure 2B. Likewise relative to African accessions, the markedly recent split time estimates between both Florida and Brazil from Gabon could reflect recent gene flow to the Americas, back migration to Africa, or shared admixture in Florida and Brazil from Gabon that predates their introduction to the Americas. Future work is needed to parse out the unique specifics of gene flow and American introductions of individual groups of samples.

We find all introduced accessions have similar levels of diversity, reduced by only ∼33-40% from the ancestral levels. Diversity in these accessions therefore remains high (*π* ≈ 0.02 at synonymous sites), reflecting a model where *N_e_* only dropped to the tens of thousands in South American populations before quickly recovering. We note that population structure in the US means the diversity estimates and measures of the efficacy of selection we report here are inflated, with underlying populations possibly exhibiting lower diversity than in South America given the recent low *N_e_* in each US accession of about 5,000 (Figure 4A). Nevertheless, diversity levels in the US when grouped into a single population are still estimated to be reduced relative to ancestral levels. We additionally find evidence of only subtle reductions in the efficacy of selection relative to the ancestral range, as reflected in an increased *π*_0_/*π*_4_ and an increase in the proportion of weakly deleterious mutations in the DFE. We do additionally find evidence that the efficacy of selection around selective sweeps is similar between African and American accessions, suggesting that the genome-wide signal of a weakly increased *π*_0_/*π*_4_ in the Americas may be due primarily to the demographic history, though future work is required to tease apart the relative contributions of demography and selection to the genomic landscape of diversity in the species, and how deleterious variation may be distributed. This genomic resilience given the introduction bottlenecks, rapid expansions, and history of eradication attempts underscores the challenge faced by ongoing and future control efforts. Despite their demography, American populations of *Ae. aegypti aegypti* all exhibit high diversity and a strong efficacy of selection, making the path of rapid adaptation to ongoing and future anthropogenic control efforts likely even in the absence of strong gene flow.

## Methods

### Data filtering

Here we make use of a previously curated and filtered whole genome sequencing dataset of 131 samples from (Love et al. 2023), which includes 27 samples from California and Florida from (Lee et al. 2019), 18 samples from Santarém, Brazil, 13 samples from Franceville, Gabon, 19 samples from Kaya Bomu, Kenya, and 20 samples from Ngoye, Senegal taken from (Rose et al. 2020), as well as 10 samples from Cali, Colombia and 24 samples from Río Claro, Colombia. Here we use the VCF from (Love et al. 2023), with detailed methods for alignment, variant calling and filtering described therein. Briefly, Colombian samples were sequenced using paired-end 150bp Illumina reads, quality checked with FASTQC 0.11.9 (Andrews 2010) and trimmed with Trimmomatic 0.39 (Bolger et al. 2014). Reads from all specimens were aligned to the AaegL5 reference genome using bwa-mem2 version 2.1 (Vasimuddin et al. 2019), then genotyped with GATK HaplotypeCaller version 4.1.9.0 (DePristo et al. 2011; Poplin et al. 2018), using the EMIT_ALL_CONFIDENT_SITES flag. SNPs were filtered using scikit-allel version 1.3.2 (DOI:10.5281/zenodo.4759368). The dataset from Love et al. (2023) that we used here imposed the following filters:

1. For the Colombian accession, specimens with mean coverage below 15X or fewer than 70% of reads mapping were removed.
2. For the full sample, indels were removed, and SNPs were filtered following GATK’s best practices (https://gatk.broadinstitute.org/hc/en-us/articles/360035890471-Hard-filtering-germline-short-variants). Specifically, sites were retained if they passed 5 of the 6 following filters: i) variant quality by depth (QD) greater than or equal to 2, ii) variant strand bias via Fisher’s exact test (FS) less than 40, iii) variant strand bias via symmetric odds ratio test (SQR) less than 4, iv) mapping quality (MQ) greater than or equal to 40, v) mapping quality rank sum test (MQRS) between -5 and 5, inclusive, and vi) site position within reads rank sum test (RPRS) between -3 and 3, inclusive.
3. All SNPs that were not called in at least 75% of specimens in at least 5 of the 6 countries were removed.
4. All non-diallelic SNPs were removed.
5. We removed regions previously identified as repetitive (Matthews et al. 2018) and regions that were not uniquely mappable.

In addition to these filters used by Love et al., for analyses that required regions of the genome not likely to have experienced strong linked or direct selection (See Population Structure and Historical effective population size inference and split times), we further filtered the genome to remove all sites within 100 kilobases (kb) of annotated genes and for locations of centromeres as reported in (Matthews et al. 2018). This mask is referred to as the intergenic sites mask in later analyses.

For diversity statistics that require an accurate count of quality invariant sites, we further removed invariant sites with more than 25% of samples missing base calls using bcftools version 1.16 (Danecek et al. 2021), as well as sites with less than a depth of 4 or greater than a depth of 30. All subsetting or filtering of VCFs for specific masks or regions was performed using bedtools v2.30.0 (Quinlan and Hall 2010) unless otherwise noted. Some analyses required sites of 0-fold or 4-fold degeneracy, which were called using a python script (https://github.com/tvkent/Degeneracy), the L5.0 version of the *Ae. aegypti* genome and annotation, NCBI accession GCF_002204515.2 (Matthews et al. 2018).

### Population structure and admixture

To infer the population structure of *Ae. aegypti,* we used two different approaches. First, we used PCA clustering as a high-level description of the underlying structure. We used the bedtools tool *intersect* to remove sites in the aforementioned intergenic sites mask from the full VCF file and converted to plink format using the --make-bed function in plink v1.90b3.45 (Purcell et al. 2007). Sites were thinned for linkage disequilibrium (LD) by removing sites in 100kb windows exceeding an r^2^ of 0.5, with a step size of 10kb. Principal components were then calculated for the full dataset and separately for samples from the Americas. Second, we estimated population structure more explicitly with ADMIXTURE (Alexander et al. 2009), using the same LD-pruned SNP set for all samples and for the Americas with k=2-10. The K number of populations with the best predictive accuracy was determined using ADMIXTURE’s cross-validation with the --cv flag, and we present results for several K around this value.

### Historical effective population size inference and split times

In order to obtain a view of the history of bottlenecks and expansions in *Ae. aegypti*, we inferred the historical effective population size of each accession. Inference of historical *N_e_* makes use of information in the site frequency spectrum (SFS) (Liu and Fu 2020a) and/or uses local sequence information to infer a distribution of coalescent times with the sequentially Markovian coalescent (SMC) (Mather et al. 2020). SMC-based methods have been shown to have the highest accuracy over times from several hundred to tens of thousands of generations ago, while SFS-based methods are most accurate in the recent past (Patton et al. 2019), assuming a large enough sample size (Terhorst and Song 2015). Because of our sample size constraints, unphased data, and the use of folded SFS’s, we use SMC++ (Terhorst et al. 2017), which has been shown to have high accuracy for a broad time frame(Terhorst et al. 2017; Patton et al. 2019), and we provide inferences from the SFS-based Stairway Plot 2 (Liu and Fu 2020a) in the supplement for qualitative assurance of size histories. For each method, we scaled generations to years and set the per generation mutation rate using estimates previously reported in (Rose et al. 2023): 0.067 years per generation and a mutation rate of 4.85 ×10^-9^.

We used SMC++ v1.15.4 (Terhorst et al. 2017), and masked the genome using our intergenic sites mask. We ran inference on all accession as described in the previous section, as well as dividing the United States into 5 putative populations determined by their clustering in PCA space and corroboration with (Lee et al. 2019): Southern California (Commerce, Mission Viejo, Garden Grove, Brawley, San Diego), Northern California (Madera, Menlo Park, Fresno), Clovis and Sanger, Exeter, and Florida (see Supplement for plots of PC’s 1-5). We converted a VCF for each chromosome to SMC++ format with the vcf2smc function in SMC++ using our mask and choosing random individuals as “distinguished” lineages. SMC++ makes use of the information captured in multiple samples by combining the coalescent times and allelic states between haplotypes of a set of one or more “distinguished” individuals, as in other SMC methods, with the allele frequencies of a set of “unlabeled samples” conditioned on the information from the distinguished individuals (Terhorst et al. 2017). For accessions with more than 5 samples, we used 5 distinguished individuals, for accessions with 3-5 samples we used 2 distinguished individuals, and for accessions with only 2 samples we chose a single distinguished individual, with the remaining individual specimens from an accession considered as unlabeled samples. We ran the estimate function in SMC++ on these data, using 10 knots, a window size of 10, and regularization penalties of 4, 5, and 6 for inference over the past 800,000 years. Because the choice of regularization penalty resulted in qualitatively different N_e_ trajectories for some accessions (see Results), we ran neutral simulations in *msprime* version 1.2.0 (Baumdicker et al. 2022) to estimate the site frequency spectra under the inferred demographic histories to independently compare the fit of SMC++ runs under the three different regularization parameters. Simulated SFSs were extracted from the simulated tree sequence using tskit version 0.5.6 (Kelleher et al. 2019) and compared to the observed SFS for each accession using a multinomial likelihood calculation following (Beichman et al. 2017); we present results for all accessions with a regularization penalty of 5, which was a better fit for our South American accessions and Gabon, and did not qualitatively alter the results for Kenya and Senegal (See Results and Supplementary Figures 8-14). We used the same parameters for the USA accessions, though we did not assess the fit of the SFS for these accessions as the sample sizes are too low to reliably use SFS-based methods and following the other accessions, the qualitative results are unlikely to be altered.

In addition to inference of *N_e_* over time, SMC++ allows for estimation of population split times via a clean split model using inferred within- and cross-coalescent rates from SMC++ inference within and between populations. To estimate split times between accessions, we used the split function in SMC++ for every accession pair. We followed the same procedure described above to determine distinguished individuals for each accession. We then converted a VCF for each chromosome to SMC++ format using the vcf2smc function, and separately for each accession using the distinguished individuals in the focal accession. We treated all remaining individuals from both accessions as undistinguished lineages. We then ran the split function in SMC++ using these joint SMC++ files and the model fits for each accession from the previous estimate step.

In addition to SMC++ *N_e_* history inference, we ran Stairway Plot 2 (Liu and Fu 2020b) on the country-level groups of accessions (splitting Colombia into Río Claro and Cali) using an intergenic sites SNP set filtered as described in Data Filtering, but masking 10kb around genes. For each accession, we calculated the folded SFS using scikit-allel (DOI:10.5281/zenodo.4759368), then ran Stairway Plot 2 using the mutation rate and generation time estimates from (Rose et al. 2023) as with SMC++, and a sequence length, *L*, of 66,563,773, representing the number of filtered variant and invariant sites used to generate the SFS.

### Diversity statistics and the distribution of fitness effects

We calculated pairwise genetic diversity as π for each accession using *pixy* (Korunes and Samuk 2021) in windows of 500kb, following (Love et al. 2023), separately for all sites, 0-fold sites, and 4-fold sites. Genome-wide averages were recalculated using the raw *pixy* output as the sum of differences (count_diffs) divided by the sum of comparisons (count_comparisons). We also report genome wide average *F_ST_* and *D_xy_* for all sites (Supplementary figures 4 and 5). Error bars for all genome-wide averages were calculated as the 95% bootstrapped confidence interval with 100 bootstraps over windows in R. *Ae. aegypti* has faced substantial eradication efforts, and as such has undergone strong, recent positive selection (Love et al. 2023), which should lead to genomic heterogeneity in the efficacy of selection due to hitchhiking (Smith and Haigh 1974). As such, we also estimated the ratio *π*_0_/*π*_4_ in regions immediately surrounding selective sweeps. For diversity statistics around selective sweeps, we filtered the Sweepfinder results from Love et al. 2023 for the top 1% of CLR windows, and calculated the statistic of interest in windows surrounding these 10kb outlier windows.

Because the behaviour of statistics like *π*_0_/*π*_4_ is unclear in nonequilibrium populations, we additionally estimated the distribution of fitness effects of new mutations (DFE) in each accession, which is thought to be more robust to nonequilibrium dynamics (Brandvain and Wright 2016). To estimate the DFE, we first calculated the folded site frequency spectrum using the sfs_folded function in scikit-allel for 0-fold, 4-fold, and intergenic sites defined as sites more than 100kb up- and downstream of annotated genes. We estimated the DFE for 0-fold degenerate sites relative to both 4-fold degenerate and intergenic sites using DFE-alpha version 2.15 (Keightley and Eyre-Walker 2007) using default parameters, and obtaining *N_e_*s bins using the prop_muts_in_s_ranges function. 95% confidence intervals were calculated by bootstrapping the site frequency spectra by site using the random.choice function in numpy (Harris et al. 2020) to randomly sample the genotype arrays and re-estimating the DFE with 1,000 replicates.

## Acknowledgements

We would like to thank members of the Matute and Schrider labs, Tom Booker, and Julia Kreiner for useful feedback and discussion. D.R.S. was supported by NIH award R35GM138286. D.R.M. was supported by NIH award R35GM148244. The funders had no role in study design, data collection and analysis, decision to publish, or preparation of the manuscript.

## Data and Code Availability

All sequencing data used in this paper are publicly available (see Love et al. 2023). All code used for the analyses presented in this paper can be found at https://github.com/tvkent/aedes_demography/.

