## Supplementary Table 1, Supplementary Figures 1-14 for "Demographic history and the efficacy of selection in the globally invasive mosquito *Aedes aegypti*"

|  | *K*=2 | *K*=3 | *K*=4 | *K*=5 | *K*=6 | *K*=7 | *K*=8 | *K*=9 | *K*=10 |
| --- | --- | --- | --- | --- | --- | --- | --- | --- | --- |
| Full | 0.20135 | 0.20645 | 0.22082 | 0.21884 | 0.22025 | 0.23524 | 0.23559 | 0.25105 | 0.2586 |
| Americas | 0.12401 | 0.12548 | 0.12822 | 0.1448 | 0.13887 | 0.14594 | 0.15338 | 0.1770 | 0.17123 |

**Supplementary Table 1:** Cross validation error for ADMIXTURE runs of *K*=2-10 for the full sample set and for the Americas samples.

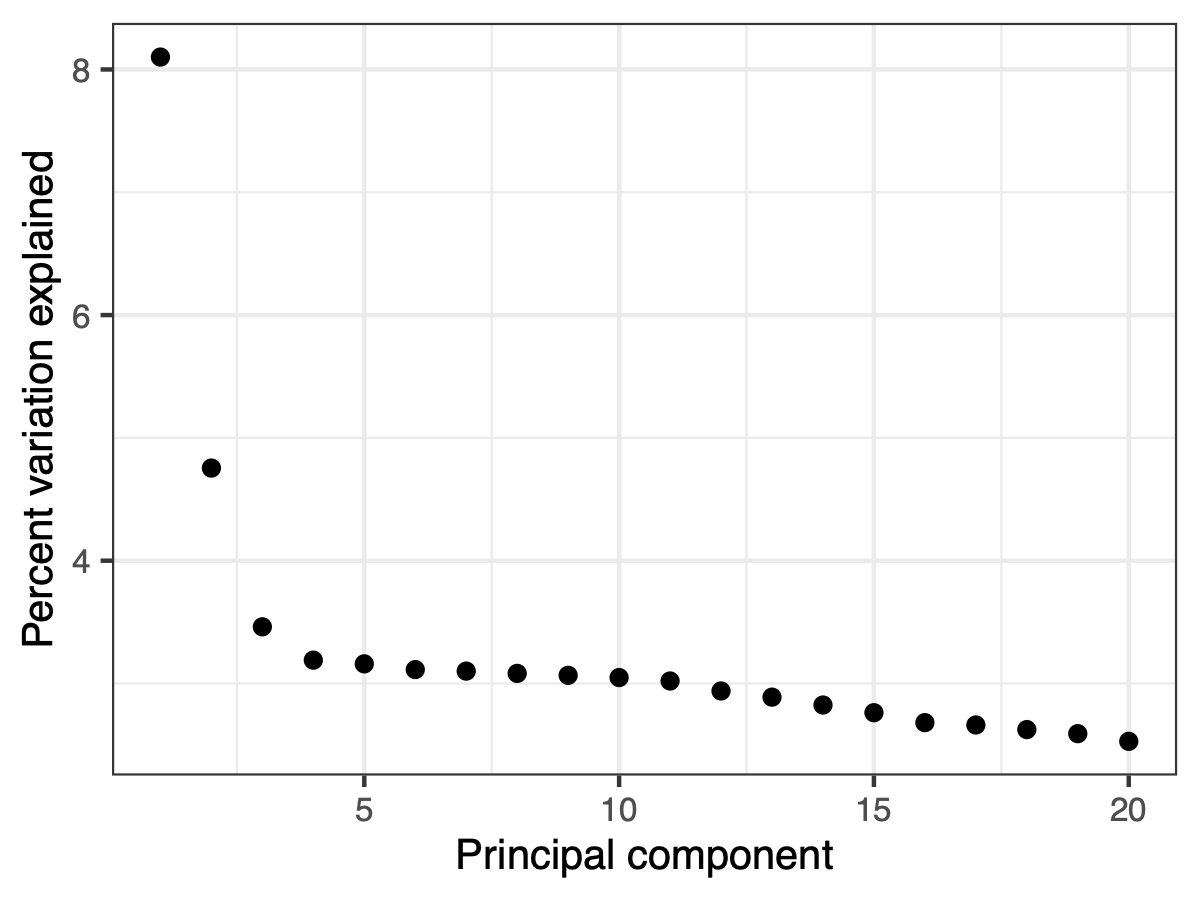

**Supplementary Figure 1:** Percent variation explained by each principal component for the country level PCA (Figure 1B).

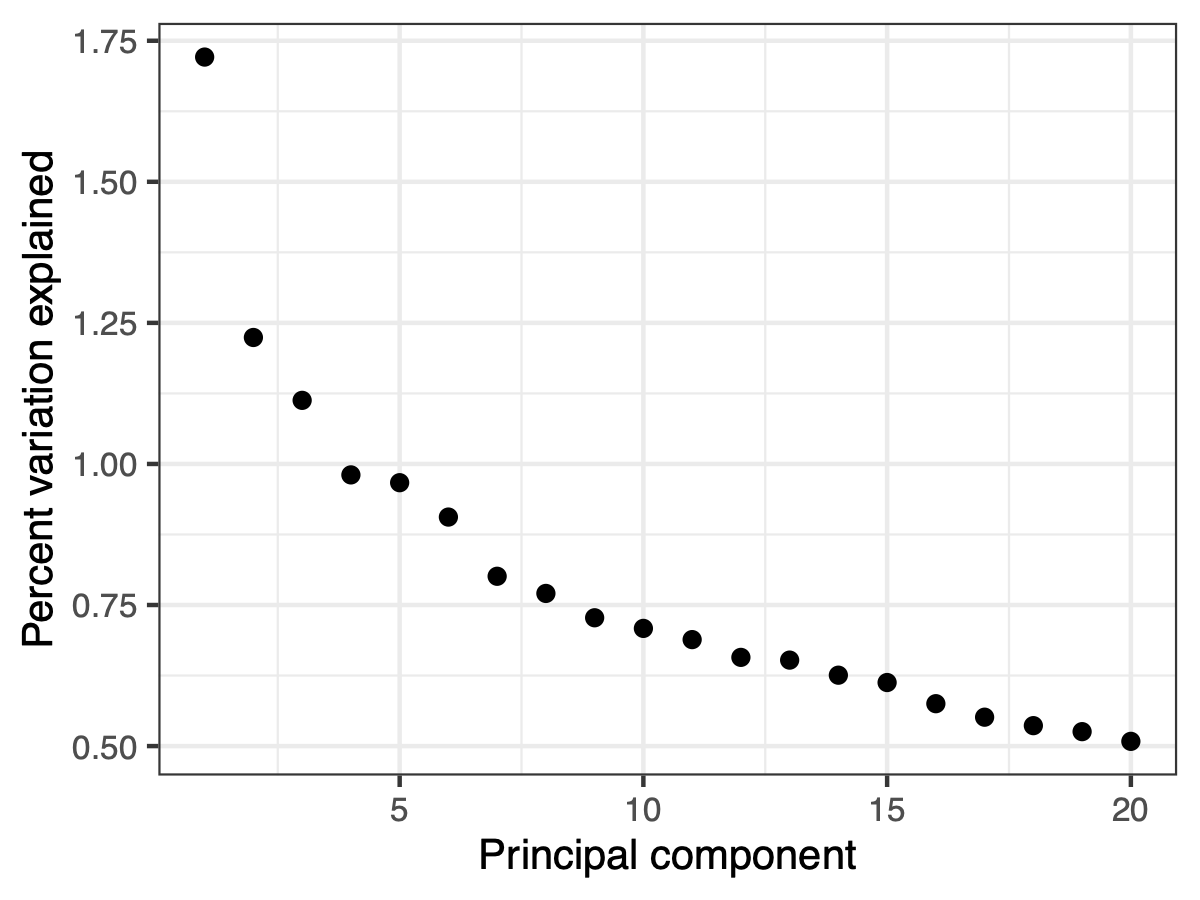

**Supplementary Figure 2:** Percent variation explained by each principal component for the country level PCA (Figure 1C).

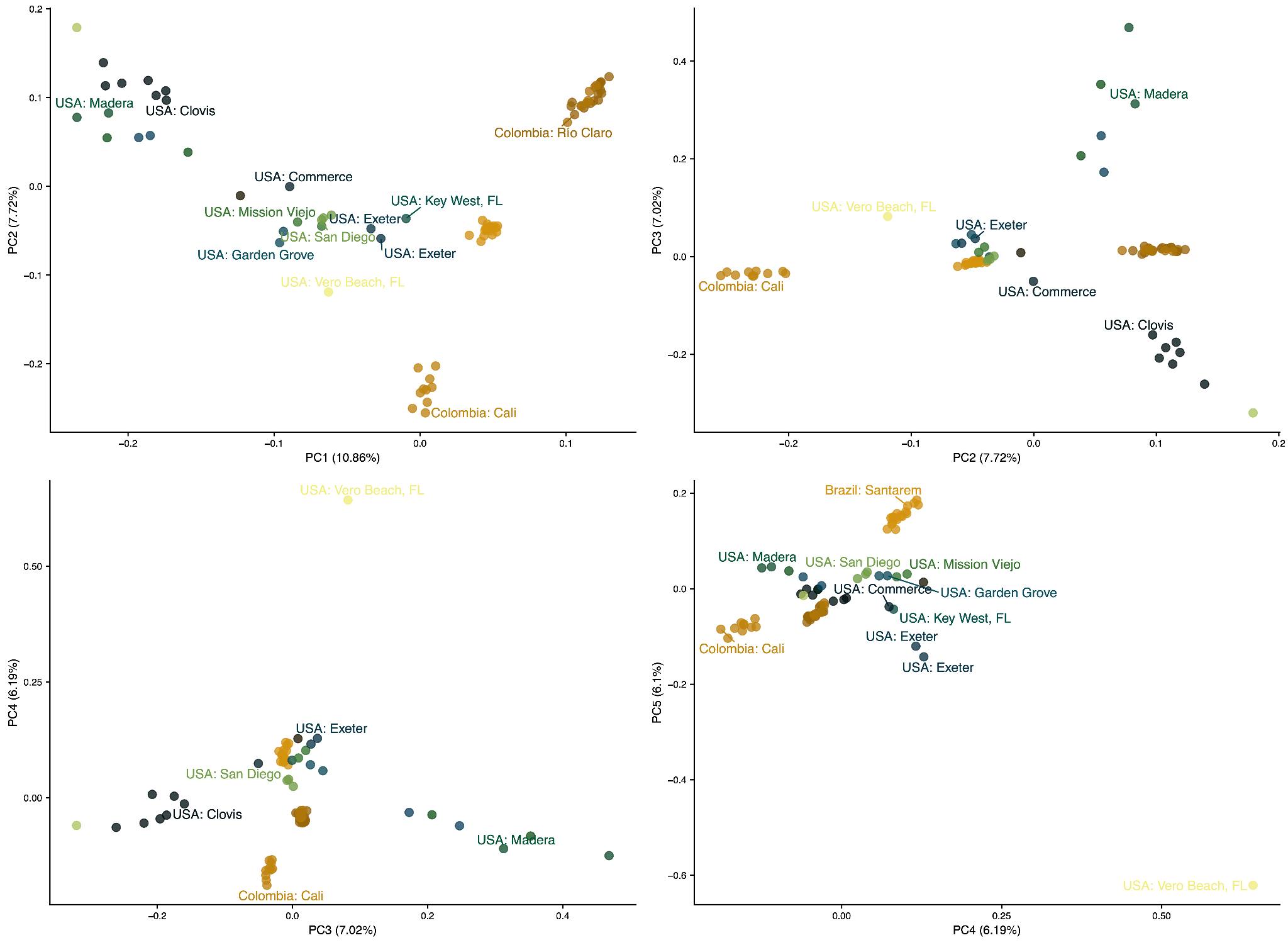

**Supplementary Figure 3:** PCA of American accessions through PC5. Major clusters in the US used in analyses are represented by the groups of Clovis and Sanger (light green), NorCal (Madera locations labeled), SoCal (Commerce and San Diego locations labeled), Florida, and Exeter.

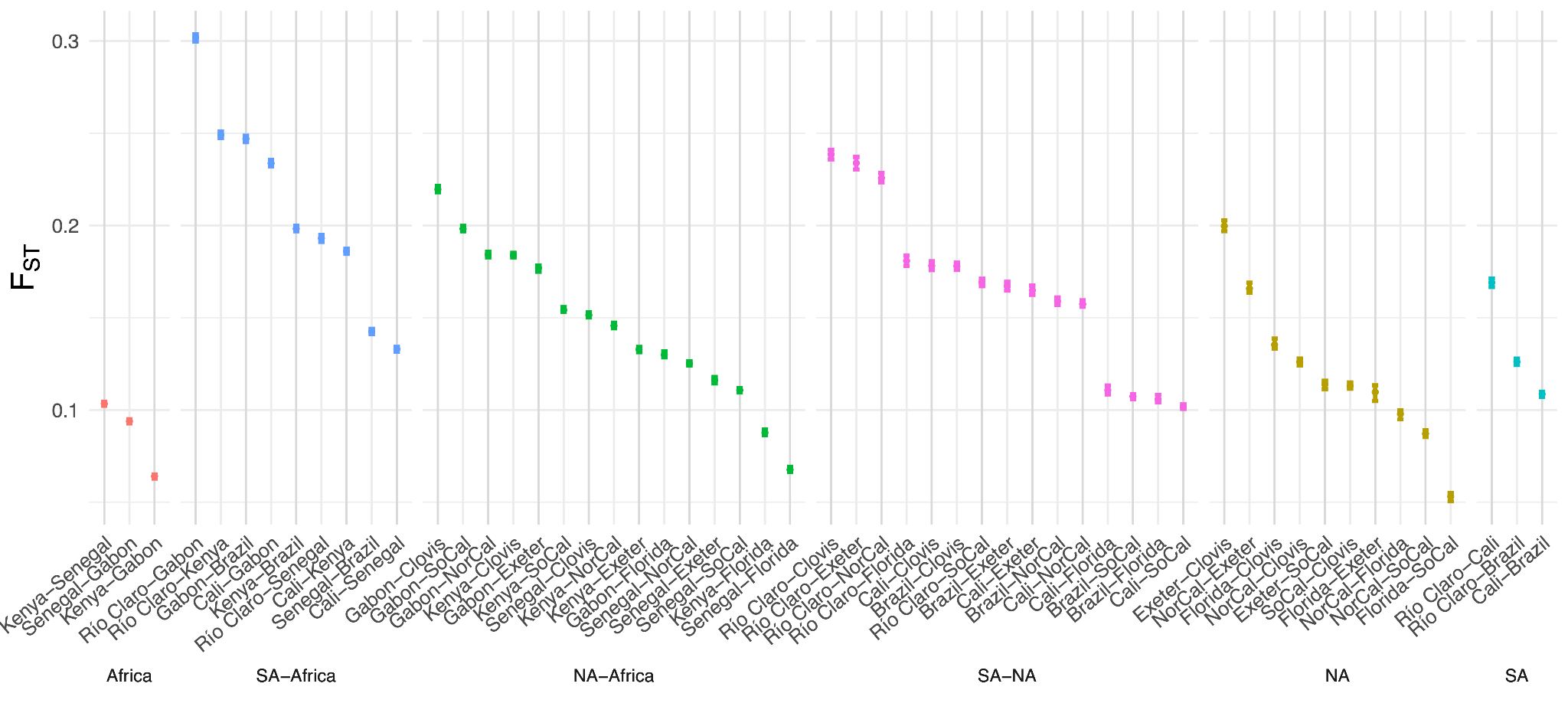

**Supplementary Figure 4:** Average *F_ST_* for all accession contrasts. NA: North America; SA: South America.

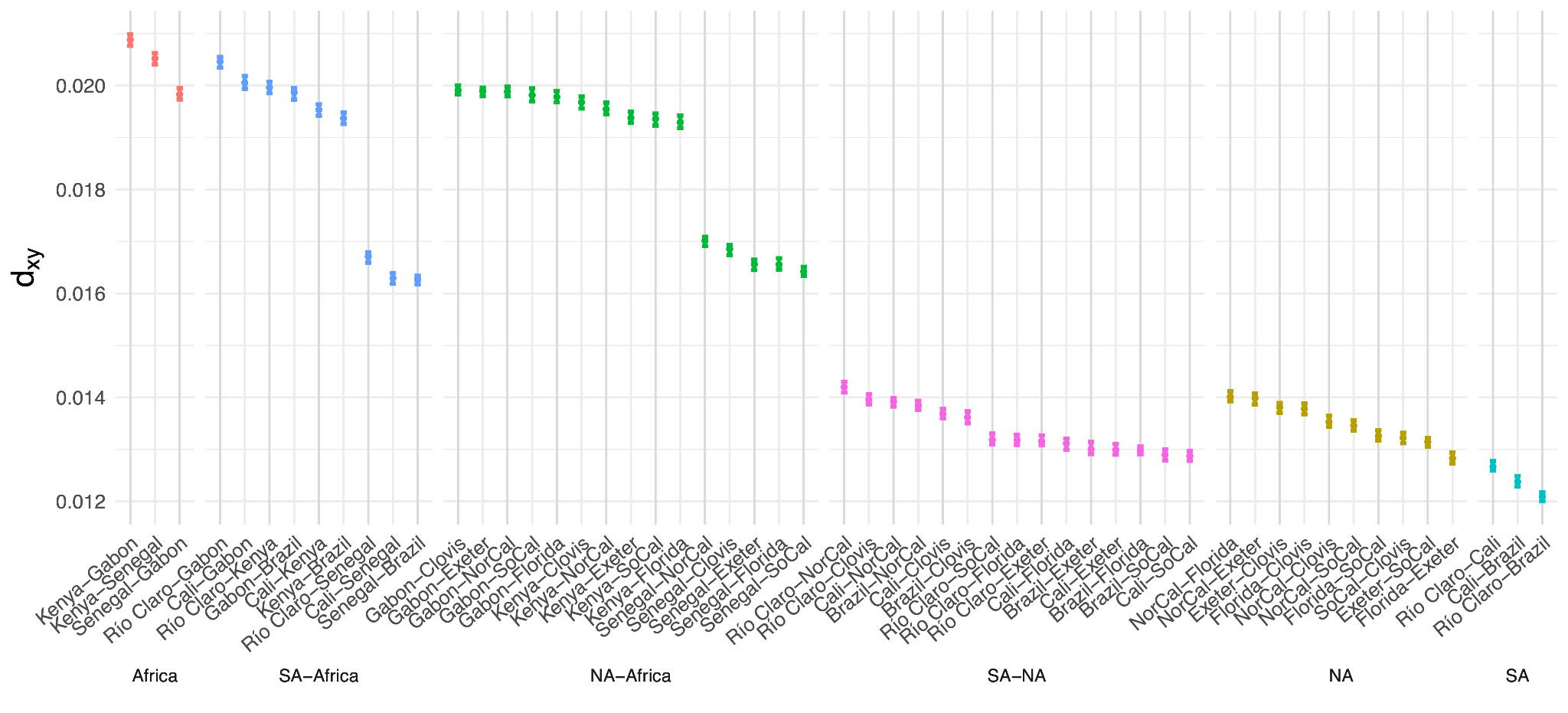

**Supplementary Figure 5:** Average *D_xy_* for all accession contrasts. NA: North America; SA: South America.

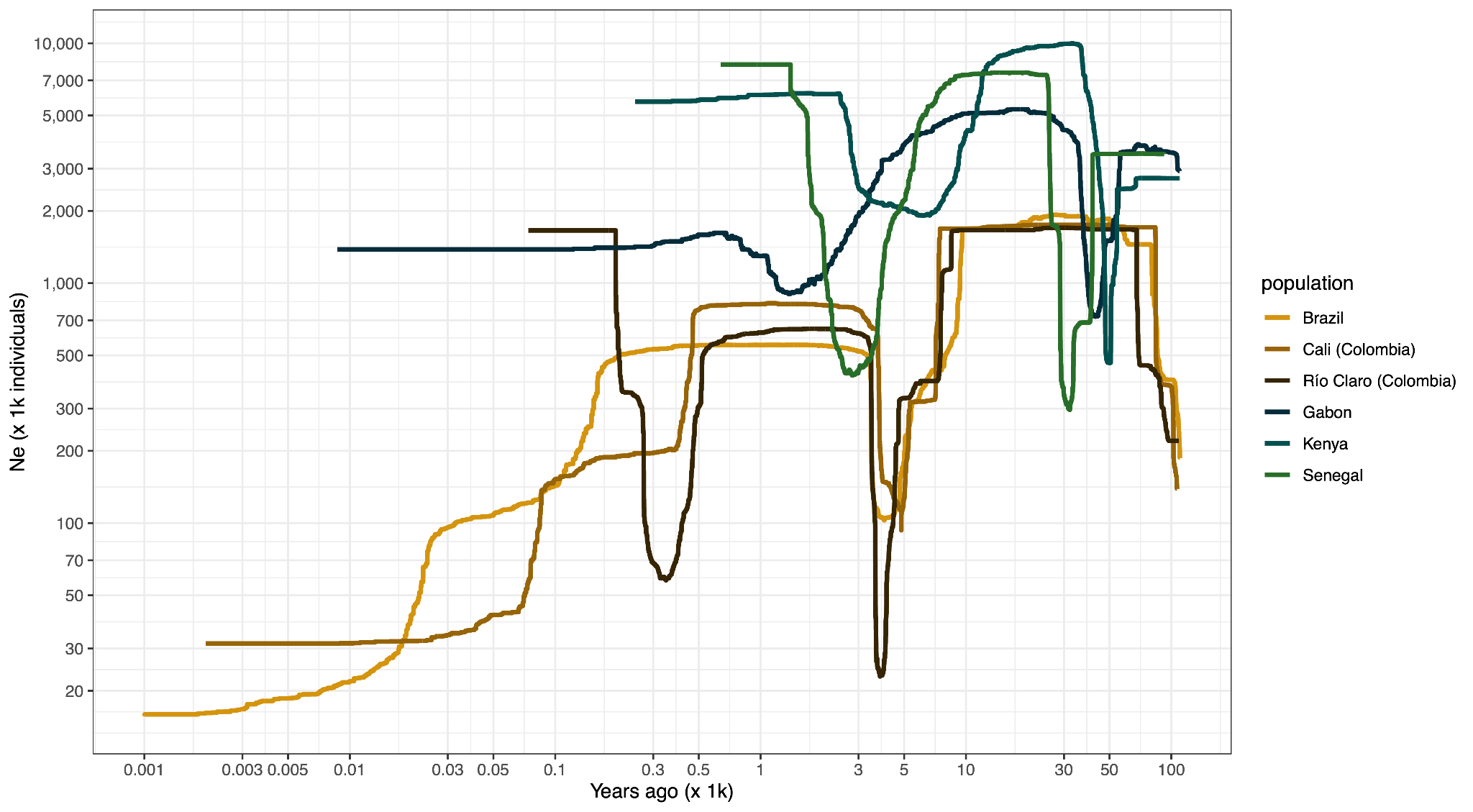

**Supplementary Figure 6:** Stairwayplot2 effective population size histories

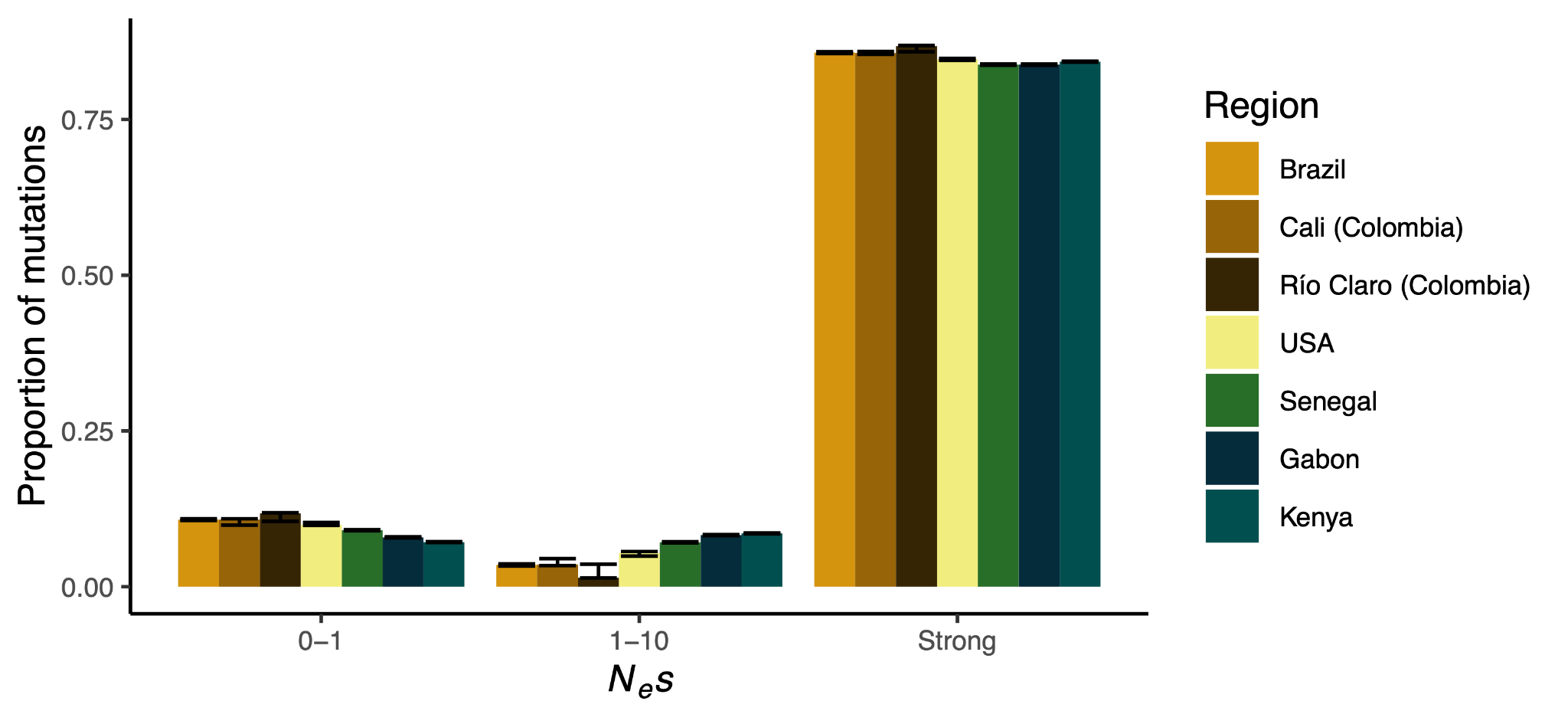

**Supplementary Figure 7:** Distribution of fitness effects combining the 10-100 and 100+ *N_e_*s bins.

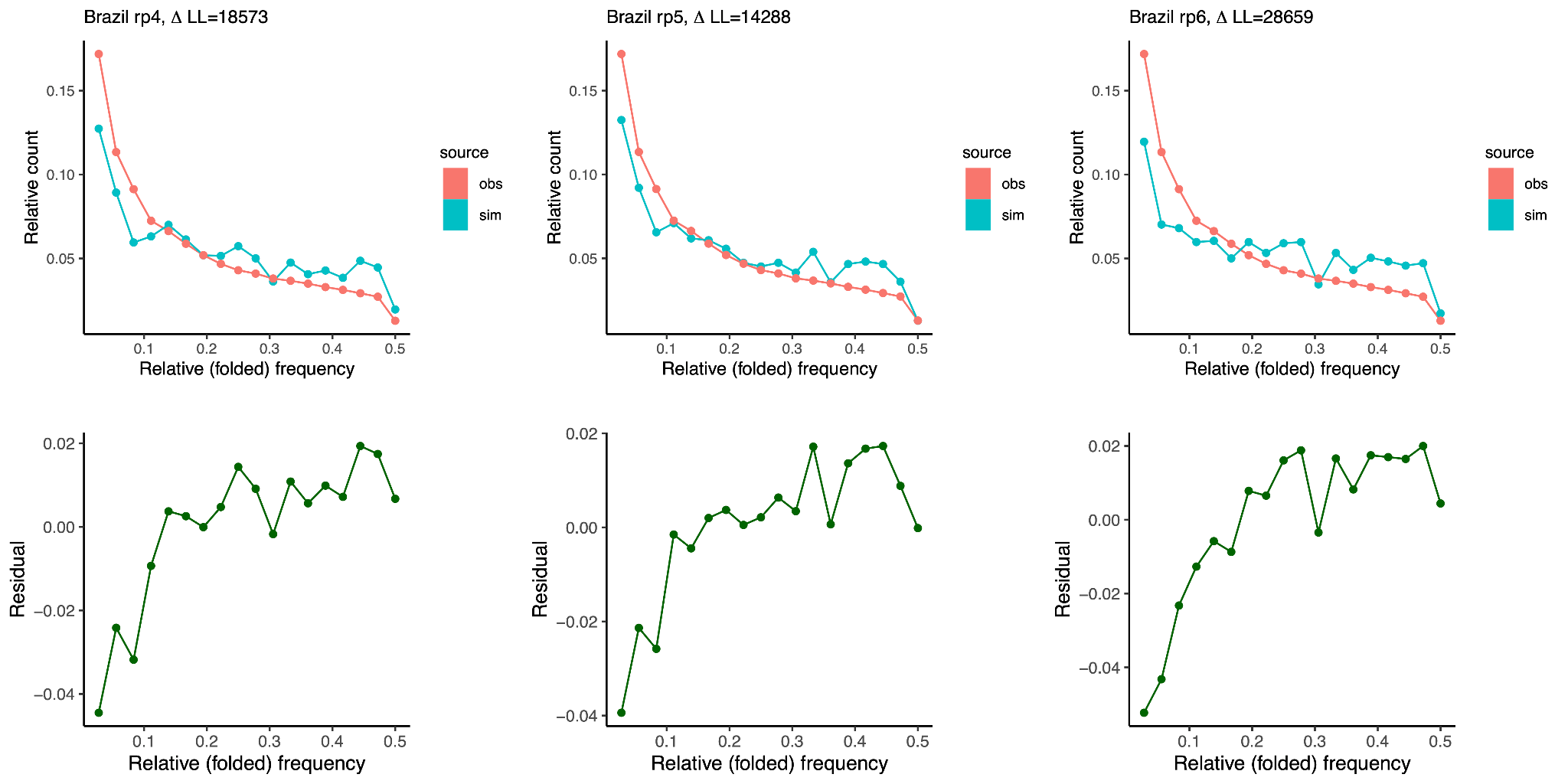

**Supplementary Figure 8:** Model fits for SMC++ regularization in Brazil. Left to right regularization parameters are 4, 5, and 6. Observed folded SFS is in red, simulated folded SFS is in blue. Residuals of the simulated SFS relative to the observed are on the bottom row.

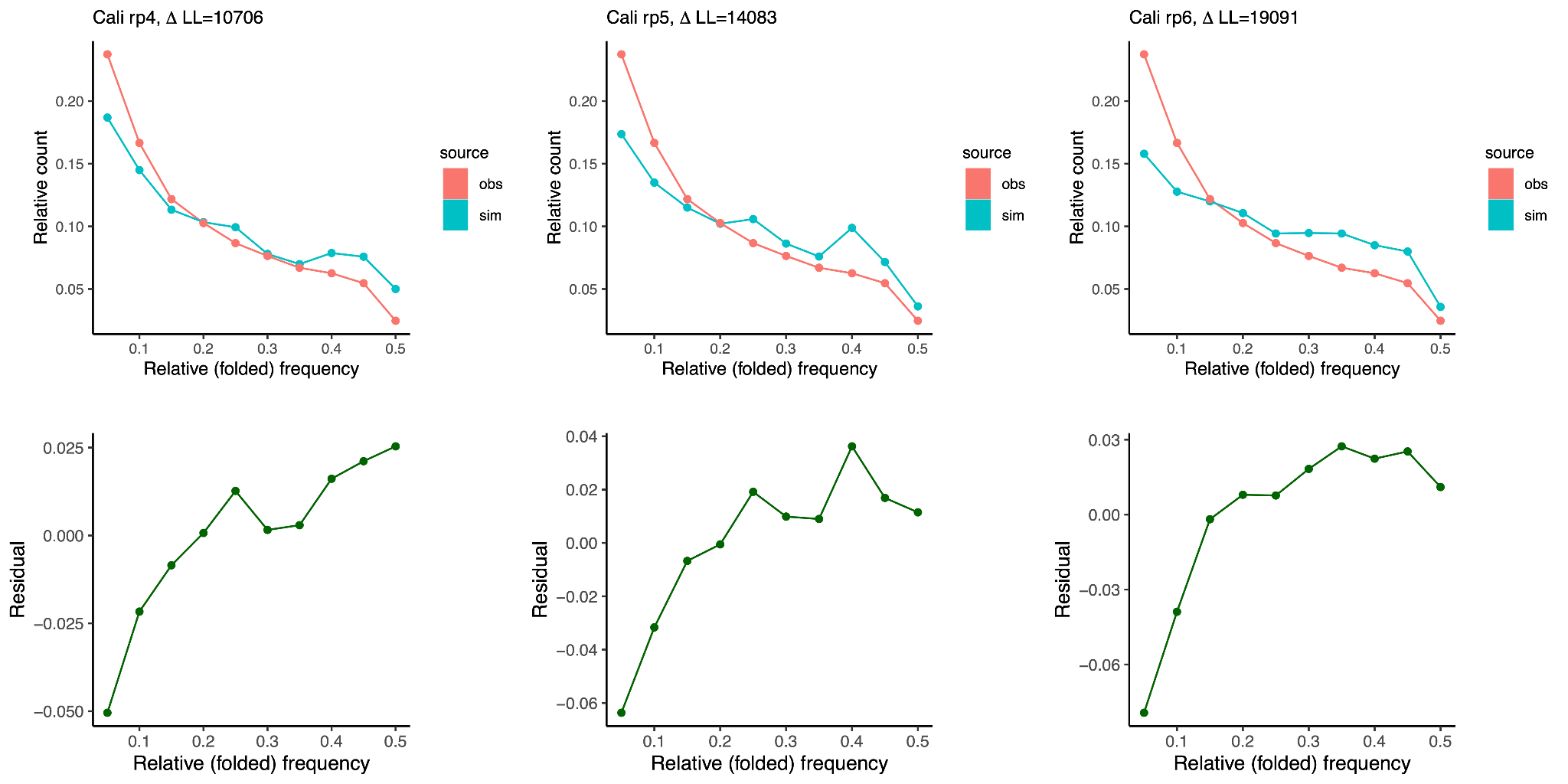

**Supplementary Figure 9:** Model fits for SMC++ regularization in Cali. Left to right regularization parameters are 4, 5, and 6. Observed folded SFS is in red, simulated folded SFS is in blue. Residuals of the simulated SFS relative to the observed are on the bottom row.

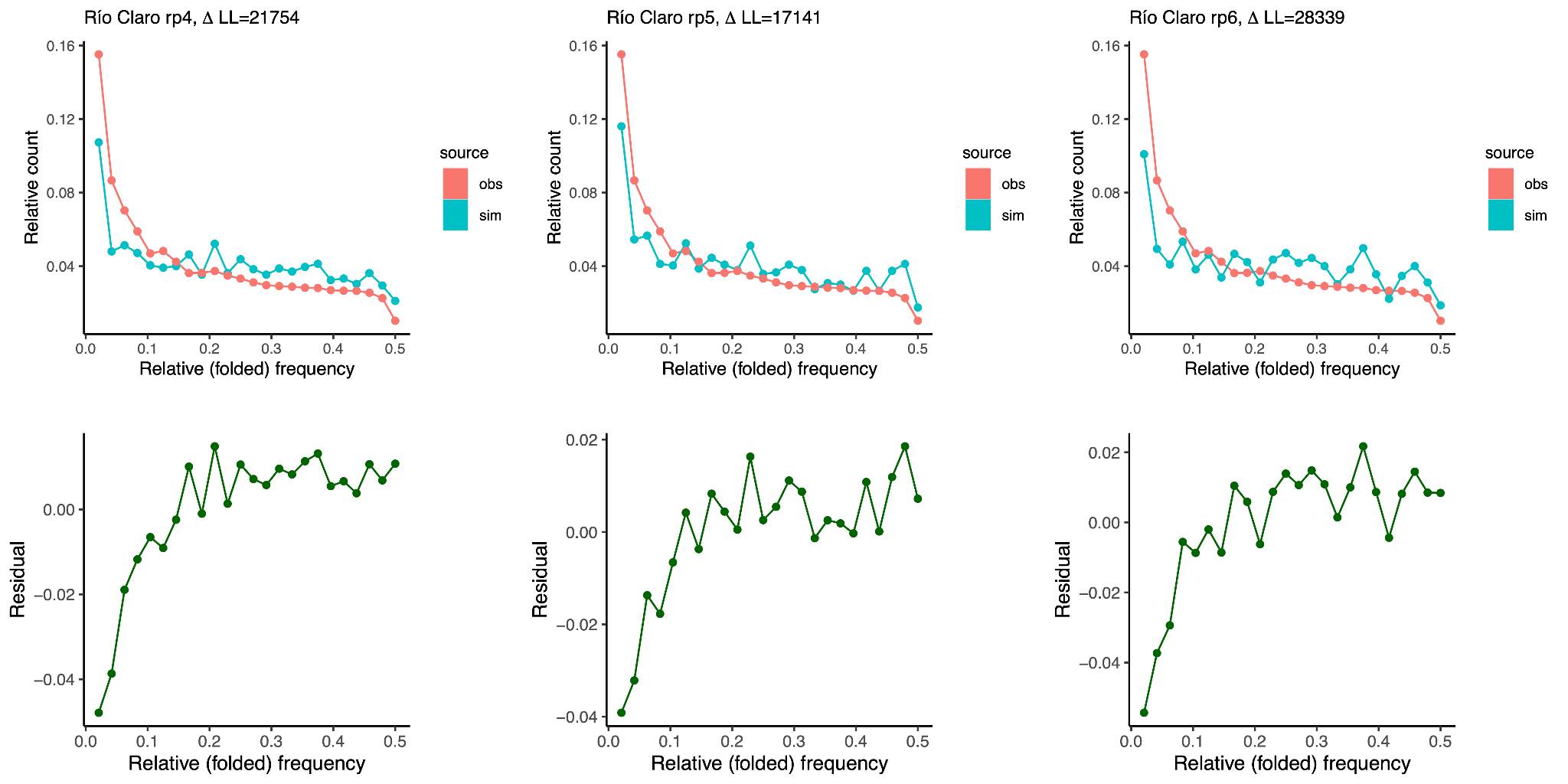

**Supplementary Figure 10:** Model fits for SMC++ regularization in Río Claro. Left to right regularization parameters are 4, 5, and 6. Observed folded SFS is in red, simulated folded SFS is in blue. Residuals of the simulated SFS relative to the observed are on the bottom row.

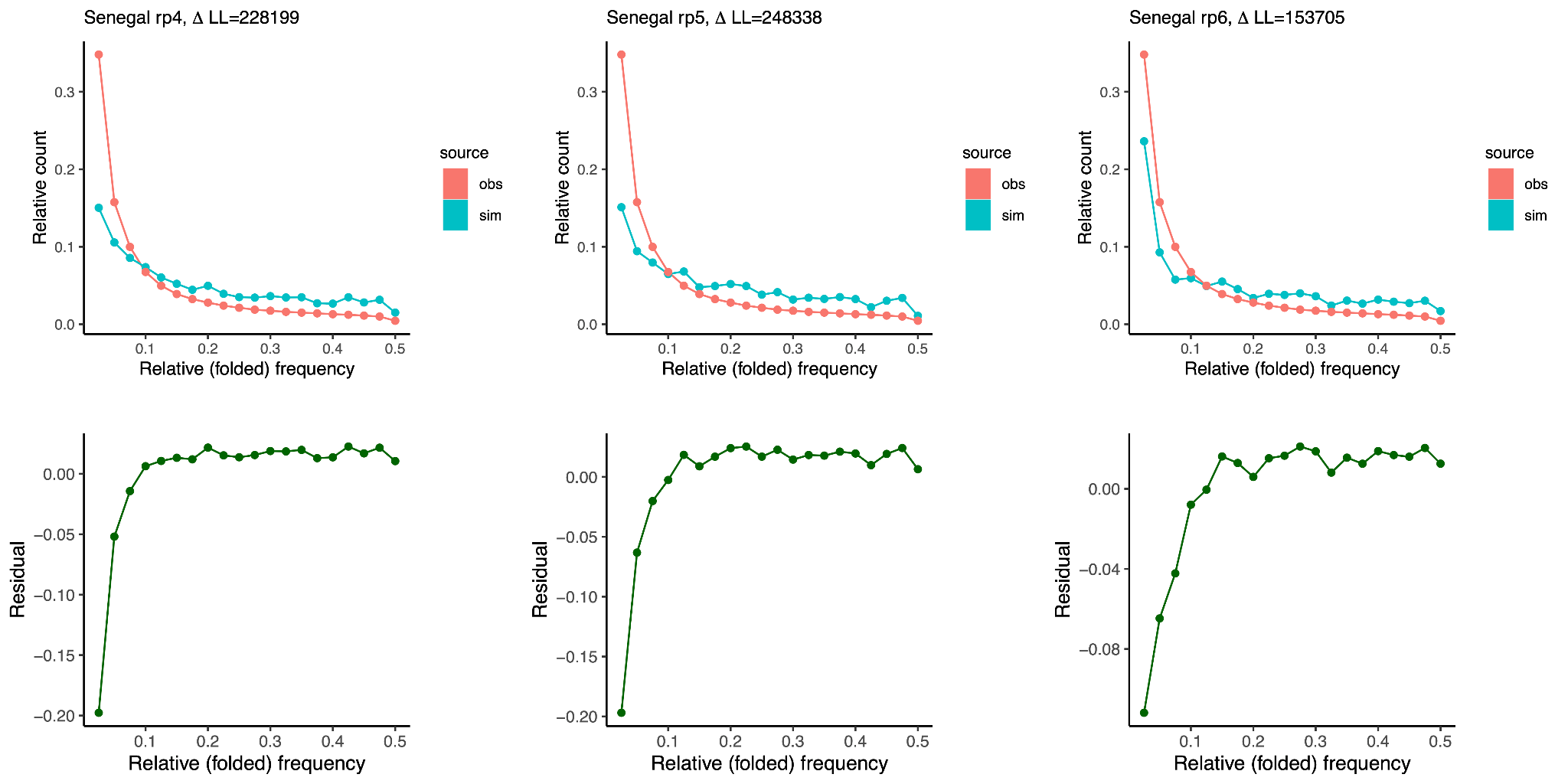

**Supplementary Figure 11:** Model fits for SMC++ regularization in Senegal. Left to right regularization parameters are 4, 5, and 6. Observed folded SFS is in red, simulated folded SFS is in blue. Residuals of the simulated SFS relative to the observed are on the bottom row.

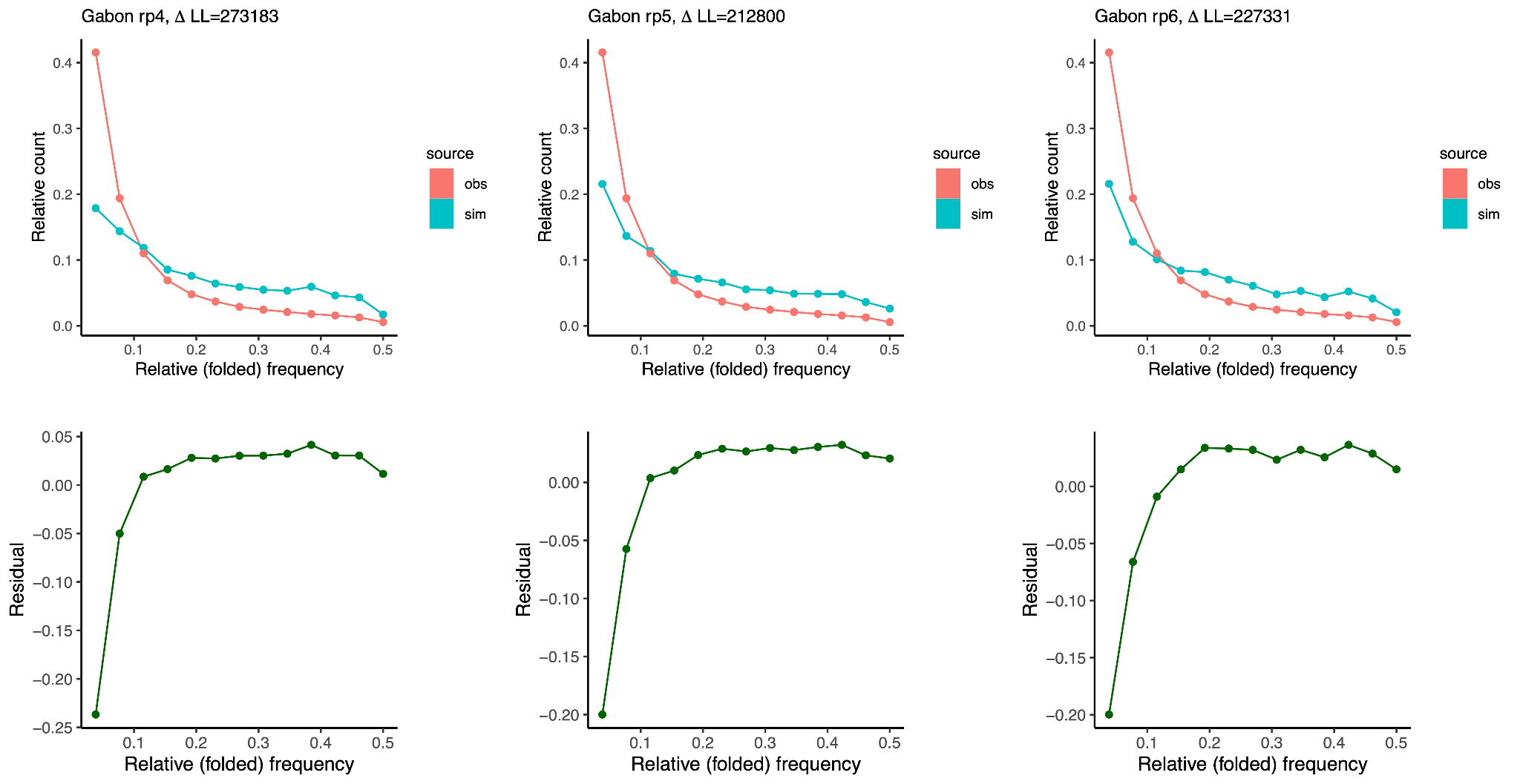

**Supplementary Figure 12:** Model fits for SMC++ regularization in Gabon. Left to right regularization parameters are 4, 5, and 6. Observed folded SFS is in red, simulated folded SFS is in blue. Residuals of the simulated SFS relative to the observed are on the bottom row.

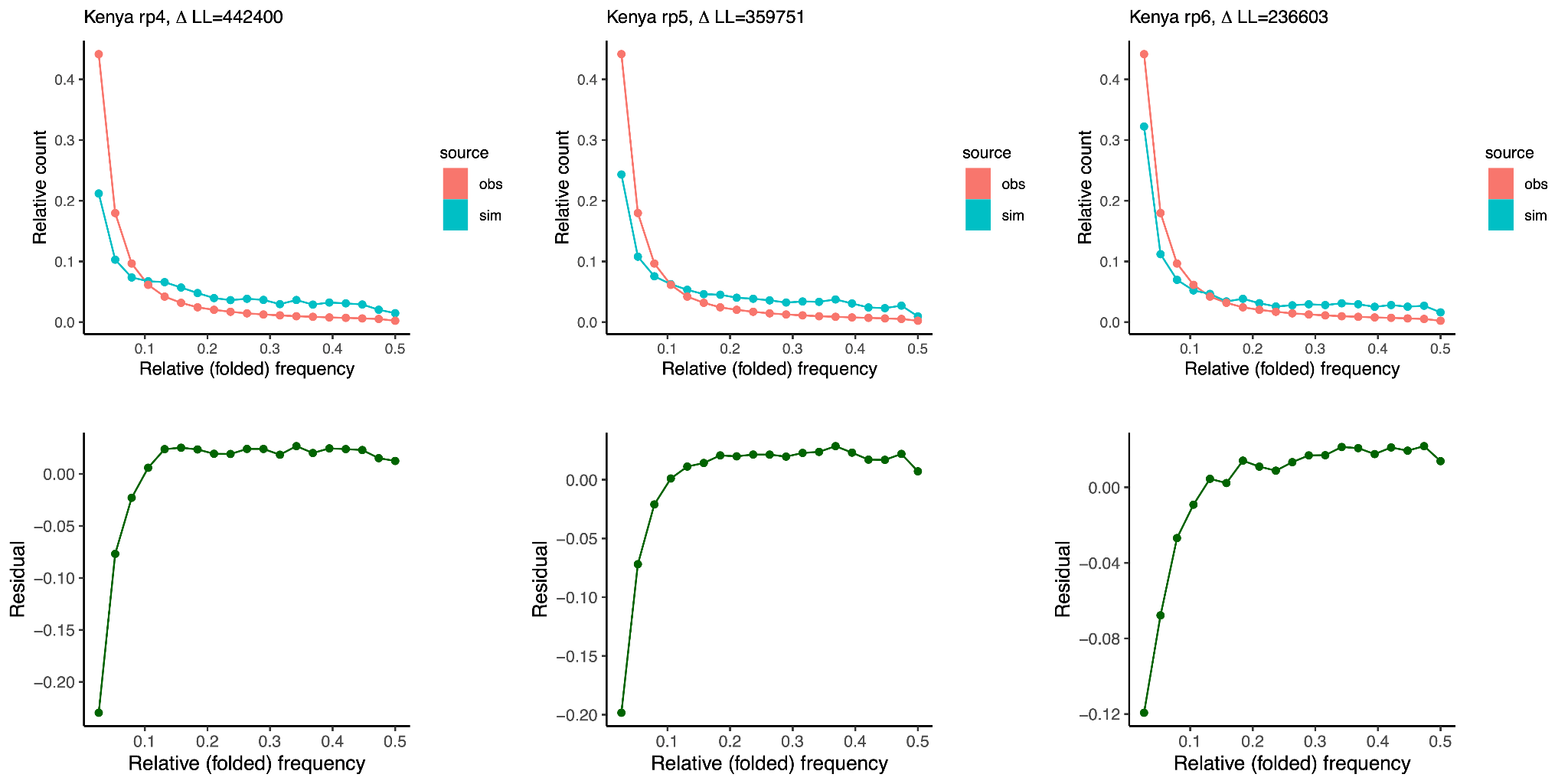

**Supplementary Figure 13:** Model fits for SMC++ regularization in Kenya. Left to right regularization parameters are 4, 5, and 6. Observed folded SFS is in red, simulated folded SFS is in blue. Residuals of the simulated SFS relative to the observed are on the bottom row.

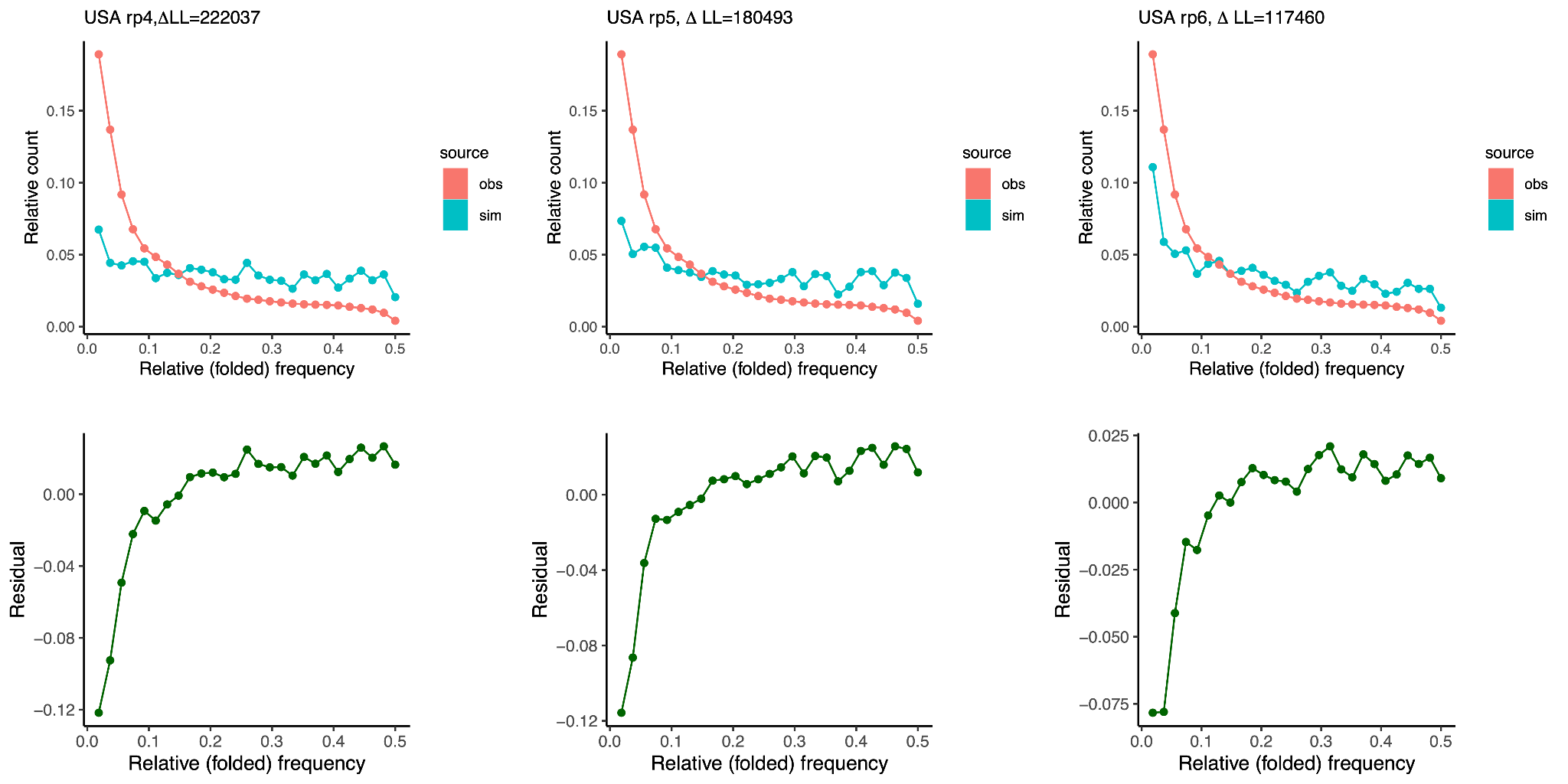

**Supplementary Figure 14:** Model fits for SMC++ regularization in the USA. Left to right regularization parameters are 4, 5, and 6. Observed folded SFS is in red, simulated folded SFS is in blue. Residuals of the simulated SFS relative to the observed are on the bottom row.
